# Seafloor topography structures subseafloor microbial communities in the Puerto Rico Trench

**DOI:** 10.64898/2026.10.13.758824

**Authors:** Corinna Breusing, William D. Orsi, Miguel Desmarais, Alvaro M. Plominsky, Irita Aylward, Dennis Graham, Robert Pockalny, Sofía Rivera Soto, Barbara Saucedo, David C. Smith, Evan A. Solomon, Jelis J. Sostre-Cortes, Arthur J. Spivack, Taylor Walton, Laura M. Wehrmann, Ying Zhang, Douglas H. Bartlett, Steven D’Hondt

## Abstract

The influence of seafloor topography on subseafloor microbial communities is poorly understood. In this study, we systematically sampled subseafloor sedimentary microbial communities and pore-water geochemistry at bathyal, abyssal, and hadal water depths of the Puerto Rico Trench. The Puerto Rico Trench exhibits vertical relief of over 8 km with seafloor topography shaped by active plate tectonics and characterized by a very steep southern wall and active mass wasting. Sampling across these features, we find that subseafloor microbial communities are influenced by topography in the following two ways. First, subseafloor sediment in the deepest part of the trench is anoxic, enriched in organic matter due to mass transport of sediment from shallower water depths, and contains an anaerobic subseafloor community that is quite distinct from the anaerobic subseafloor community in anoxic sediment at bathyal depths. The difference between these bathyal and hadal subseafloor anaerobic communities is consistent with influence of water-depth-related physical properties on subseafloor community composition. Second, subseafloor sediment remains oxygenated for meters below seafloor on steep slopes within the bathyal zone, an abyssal topographic high within the trench, and the abyssal plain seaward of the trench subseafloor, likely due to limited sediment accumulation, which selects for the presence of aerobic microbial communities. The difference between these bathyal and abyssal aerobic subseafloor communities and the anaerobic communities at bathyal and hadal depths reflects the role of seafloor topography in structuring subseafloor availability of redox reactants, which in turn drives the activities and composition of subseafloor microbial communities.

**Significance Statement:** Energy limitation is a major factor structuring microbial communities that persist deep below the seafloor. Seafloor topography has the potential to shape energy sources available to the subseafloor biosphere, but this relationship remains poorly understood. We addressed this topic by systematically analyzing subseafloor communities across depth zones and spatial features of the Puerto Rico Trench down to 20 meters below the seafloor. The diversity and metabolic potential of subseafloor communities were indirectly structured by seafloor topography due to its control on sediment accumulation rates, porewater geochemical profiles, and the availability of high-energy terminal electron acceptors for respiration. These findings provide new insights into how seafloor topography can structure microbial communities in the deep biosphere.

## Introduction

The hadal zone, which encompasses depths greater than 6000 m, is the least explored oceanic habitat on Earth (1, 2). This zone is primarily composed of topographically isolated trenches that form at subduction zones (3). Organisms living in hadal trenches need to cope with a variety of challenging conditions, first and foremost extreme hydrostatic pressure, which can reach over 1000 times the amount at sea level (1). Trenches are characterized by high sediment accumulation rates, due to pulses of mass sediment deposition that result from turbidity currents triggered by earthquakes in these tectonically active regions (4). Multiple pioneering studies in the Pacific Ocean have highlighted trenches as sedimentary depocenters with elevated levels of organic matter and microbial activity that are distinguished from other productive marine sediments by impacts from these mass deposition events and low bioturbation (5–11). Within the trench, dissolved oxygen and organic matter are rapidly consumed by aerobic microbes in the upper sediment layers, resulting in undetectable oxygen levels a few centimeters or decimeters beneath the seafloor and distinct redox stratification in deeper subsurface sediment (12).

Previous studies have shown that deep-sea sedimentary communities are structured across trench subseafloor redox zones due to selective pressures and dispersal limitations (13–17) but are also shaped by geographic location (14, 17, 18) and water depth (17, 19–21).

In addition, seafloor topography has emerged as an important factor influencing the distribution and activity of microbial communities living in seafloor surface sediment of deep-sea trenches (13, 15, 21–23). However, whether seafloor topographic features also impact the microbial communities persisting deeper below the seafloor is unknown as obtaining long sediment cores (>10 meters) suitable for microbiological study is technically challenging in the deep sea, especially at hadal depths (24, 25).

With a vertical relief characterized by steep walls, topographic highs and water depths exceeding 8,300 m, the Puerto Rico Trench represents an ideal system to assess the impact of seafloor topography on subseafloor microbial communities. The Puerto Rico Trench contains the deepest point in the Atlantic Ocean, which is formed by a combination of strike-slip movements, tectonic deformation and oblique subduction (26), resulting in a complex, asymmetrical V-shaped topography. Furthermore, a near-seafloor unconformity occurs along the trench slope, which allows a greater depth of oxygen penetration into the subsurface sediment (27), thus likely affecting the composition and metabolic functions of associated microbial communities.

In this study, we embarked on a deep-sea long coring expedition to investigate potential effects of seafloor topography on the deep biosphere within the Puerto Rico Trench region. We systematically sampled sediment communities from unique seafloor topographic features of the trench across bathyal, abyssal and hadal depths (Table S1; Fig. 1). Our dataset includes samples from a broad range of sediment depths, extending down to 20 meters below seafloor (mbsf). For context, our dataset includes comparative seawater samples from the same sites. Synthesis of geochemical, metagenomic and 16S rRNA barcoding datasets from this unique deep-sea sediment core collection allowed us to assess how seafloor topography structures the availability of energy sources and the composition of subseafloor microbial communities.

**Fig. 1.**
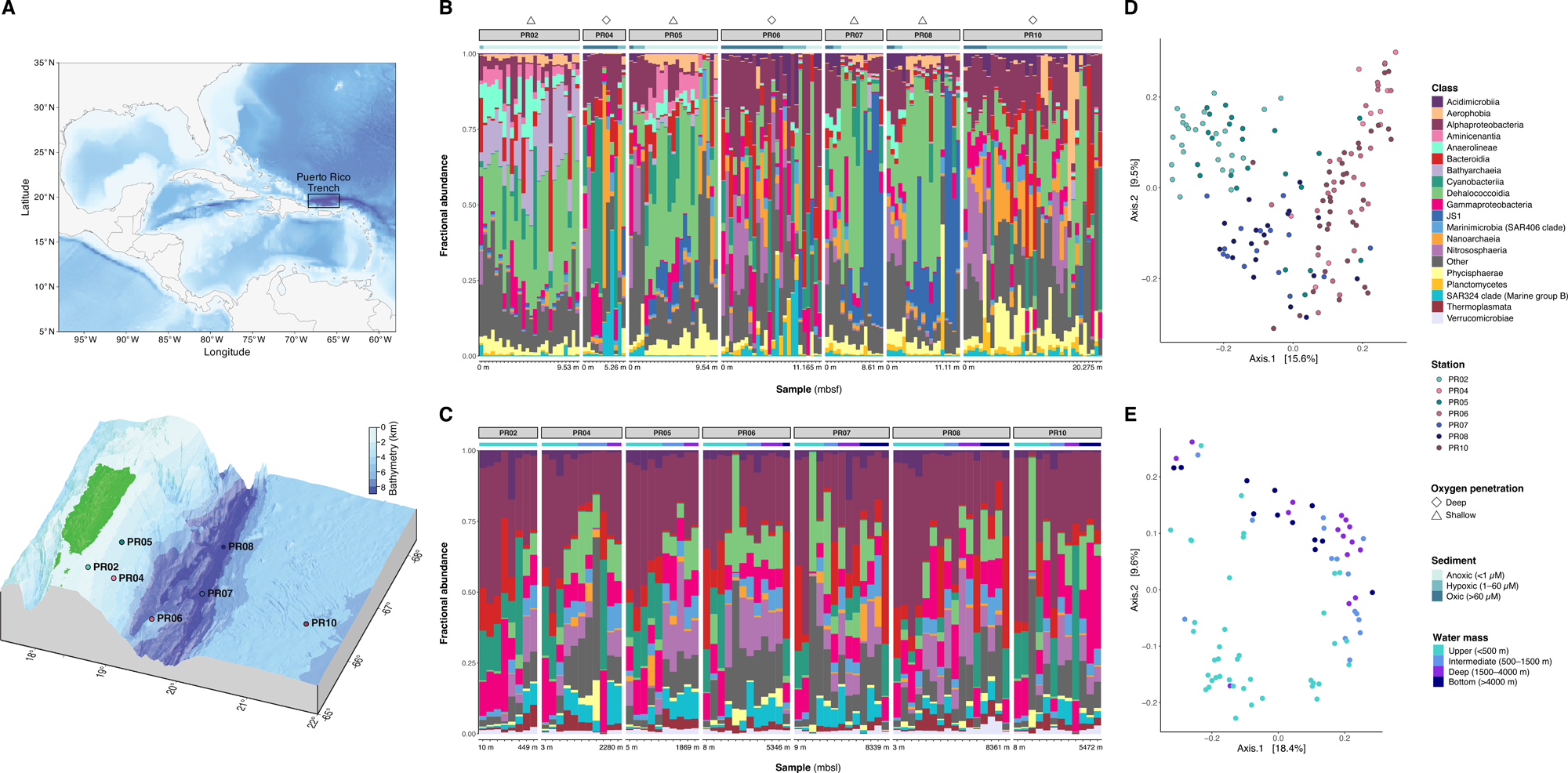
Map of the seven sampling stations (A) and fractional abundances of microbial phyla in sediment (B) and seawater (C) samples based on 16S rRNA amplicon analyses. Sediment samples are ordered by meters below seafloor (mbsf) and annotated with the respective oxygen level. Oxygen penetration at the corresponding station is shown with triangles (for low penetration) and diamonds (for high penetration). Seawater samples are ordered by meters below sea level (mbsl) and annotated with the respective water layer from which the sample was collected from. Water layers were classified based on ref. (80). Sediment samples show differences in taxonomic composition related to station and oxygen penetration level (B) that result in three distinct clusters of microbial communities in principal coordinate analyses based on unweighted UniFrac distances (D): 1) bathyal (shelf and upper slope) communities with low oxygen penetration at the sampling station (PR02, PR05), 2) hadal communities with low oxygen penetration at the sampling station (PR07, PR08) and 3) bathyal to abyssal communities with deep oxygen penetration at the sampling station (PR04, PR06, PR10). Seawater samples do not show any observable clustering by station but are structured by water depth (E).

## Results

### Topographic features of the Puerto Rico Trench

The Puerto Rico Trench is a deep, narrow, steep-walled submarine feature with ca. 8 km vertical relief (Fig. 1A). The trench floor contains both extreme topographic lows and local topographic highs. We cored two hadal sites in the bottom of the trench corresponding to the deepest part of the Atlantic Ocean (PR07 at 8359 mbsl, PR08 at 8385 mbsl). To characterize relationships between trench topography and subseafloor communities, we also cored an upper bathyal site on the Puerto Rico shelf (PR02 at 493 mbsl), two bathyal sites on the slope connecting the shelf to the trench (PR04 at 2598 mbsl, PR05 at 1902 mbsl), an abyssal site on a topographic high within the trench (PR06 at 5373 mbsl), and an abyssal plain site seaward of the trench (PR10 at 5431 mbsl).

### Porewater and sediment chemistry

Dissolved oxygen (O_2_) declines rapidly with sediment depth at the upper bathyal stations (PR02, PR05) and the hadal stations (PR07, PR08), leading to anoxia (<1 µM O_2_) a few centimeters to decimeters beneath the seafloor (Fig. S1). By contrast, at the lower bathyal site (PR04) and the abyssal sites (PR06, PR10), dissolved O_2_ penetrates deeply and the sediment is oxic (>60 µM) to hypoxic (1–60 µM) meters beneath the seafloor (Fig. S1). Dissolved nitrate concentrations generally decrease with depth below seafloor at all stations, with strongest declines at the upper bathyal sites (PR02, PR05) and the hadal sites (PR07, PR08) (Fig. S2). Dissolved sulfate concentrations decrease strongly with sediment depth at the hadal stations (PR07, PR08) but they are relatively constant throughout the cored interval at the bathyal and abyssal sites (Fig. S3).

Alkalinity profiles provide evidence of respiration at all sites, increasing rapidly with sediment depth at the hadal stations (PR07, PR08) and slightly with sediment depth at the bathyal and abyssal sites (Fig. S4).

Sediment chemistry also shows notable differences between stations related to seafloor topography (Fig. S5–S8; Supplementary Results), with organic matter concentrations being overall highest on the bathyal shelf site (PR02) and the two hadal stations at the trench bottom (PR07, PR08). The hadal stations exhibit the most complex profiles, including intervals of elevated calcium carbonate, total organic carbon (TOC) and total organic nitrogen (TN) content as well as reversals in δ^13^C_org_ values indicative of past mass wasting events. Bulk adenylate concentrations (a proxy for microbial biomass) decrease with sediment depth at all stations (Fig. S9). This decrease is weakest at the hadal sites (PR07, PR08) and most pronounced at the abyssal sites (PR06, PR10) and the bathyal site (PR04), where subseafloor adenylate concentrations drop by three orders of magnitude in the first meter below the seafloor (Fig. S9).

### Microbial community composition and structure

16S V4-V5 metabarcoding in 166 sediment samples and 105 comparative seawater samples recovered 22,859 final amplicon sequence variants (ASVs) spanning at least 74 microbial phyla. Compositions of the sedimentary and seawater microbiomes based on these ASVs are overlapping yet distinct (Fig. 1, 2, S10; Table S2, S3; PERMANOVA: p < 0.004, R^2^ = 0.0091– 0.0571). The sedimentary communities are overall dominated by taxa from the phyla Chloroflexi (mean: 23.30%, mostly Dehalococcoidia), followed by Proteobacteria (mean: 17.30%, mostly Alpha- and Gammaproteobacteria), Crenarchaeota (mean: 7.95%, mostly Nitrososphaeria and Bathyarchaeia), Bacteroidota (mean: 6.13%, mostly Bacteroidia), Caldatribacteriota (mean: 6.13%, mostly JS1), Planctomycetota (mean: 5.95%, mostly Phycisphaerae), and Cyanobacteria (5.64%, mostly Cyanobacteriia) (Table S2; Fig. 1, S11). Taxon abundances varied from site to site related to station (i.e., seafloor topography), sediment depth and the corresponding geochemistry, with oxic/hypoxic communities being distinct from anoxic communities (Fig. 1, 2, S11; Table S3; PERMANOVA: p = 0.001, R^2^ = 0.0216–0.1452). In total, three distinct community clusters are observed: 1) bathyal (shelf and upper slope) sites with low oxygen penetration (PR02, PR05), 2) hadal sites with low oxygen penetration (PR07, PR08) and 3) bathyal and abyssal sites with deep oxygen penetration (PR04, PR06, PR10). The strongest compositional differences separate stations on the shelf and upper slope from stations on the slope, mid-trench high and abyssal plain. For example, Dehalococcoidia and Anaearolineae are prominent at all stations with little oxygen penetration (PR02, PR05, PR07, PR08), whereas Nanoarchaeia, Nitrososphaeria and Alphaproteobacteria are overall more abundant at stations with deep oxygen penetration (PR04, PR06, PR10). In addition, Aminicenantia are most frequent at the bathyal, anoxic stations (PR02, PR05), JS1-group Caldatribacteriota are dominant in deeper, anoxic hadal sediments (>2 mbsf) and Bathyarchaia are more adundant at shallower depths (Fig. 1, S11).

**Fig. 2.**
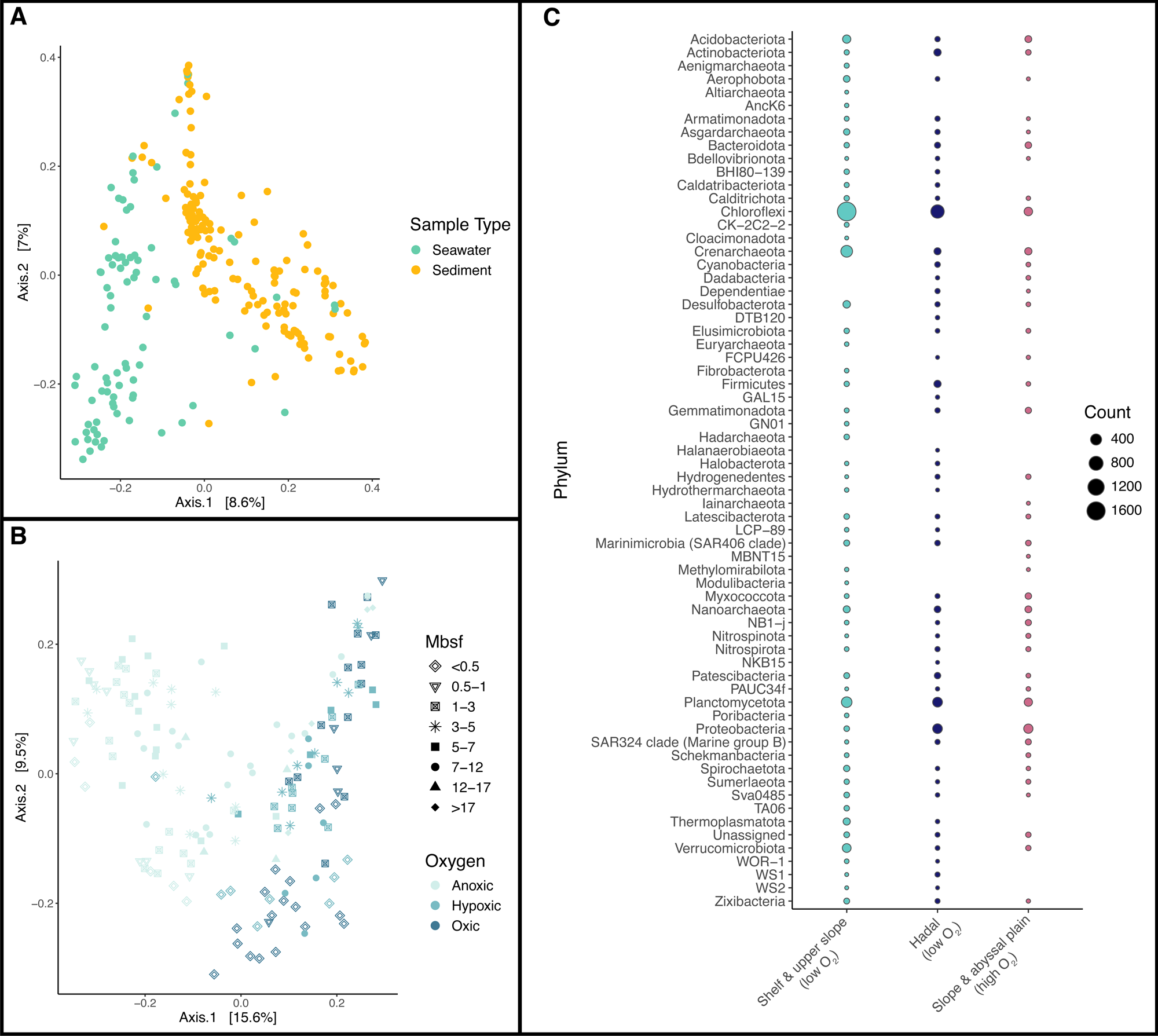
Structuring of microbial communities by habitat type (A), partitioning of sedimentary microbiomes by subseafloor oxygen availability (B) and differential abundance of ASVs between community clusters defined in Fig. 1 (C). Despite overlap in microbial taxa, sediment and seawater habitats contain distinct communities. Sediment communities are shaped by oxygen concentration, with 3,726 ASVs being enriched at the bathyal sites with low oxygen penetration, 1,916 ASVs being enriched at the hadal sites with low oxygen penetration, and 1,169 ASVs being enriched at the slope and abyssal plain sites with deep oxygen penetration. Only ASVs that are overrepresented in one cluster compared to all others are shown. Principal coordinate plots are based on unweighted UniFrac distances.

Seawater communities share most of the dominant phyla with sediment communities, but in different relative proportions, with Proteobacteria being most abundant (mean: 41.20%), followed by Chloroflexi (mean: 9.50%), Crenarchaeota (mean: 8.29%), Bacteroidota (mean: 7.82%), and Cyanobacteria (mean: 7.36%) (Table S2; Fig. 1, S12). Seawater samples also exhibit higher average proportions of Marinimicrobia, SAR324 clade bacteria and Thermoplasmatota compared to sediment samples and seem to be primarily shaped by water layer (Fig. 1, S11; Table S3; PERMANOVA: p = 0.001, R^2^ = 0.0985–0.1597). Although differences in community composition are most evident between the upper 500 m and all remaining intervals >500 m, structuring is also observed between the intermediate (500–1500 m), deep (1500–4000 m) and bottom (>4000 m) water layers (Fig. 1; Supplementary Discussion).

### Metabolic potential of sediment communities based on metagenome-assembled genomes (MAGs)

2,395 moderate- to high-quality MAGs were recovered across 80 sediment samples, including 827 MAGs that are nearly complete (>90% completeness, <5% contamination) (Fig. 3A; Table S4). 1,781 MAGs (74.36%) could be assigned to genus, while 616 MAGs (25.72%) are classified to species with a representative in the Genome Taxonomy Database. Using a dereplication threshold of 95% ANI, 1,393 out of the 2,395 MAGs (58.16%) represent unique species, although actual species diversity might be higher given that MAGs are incomplete.

**Fig. 3.**
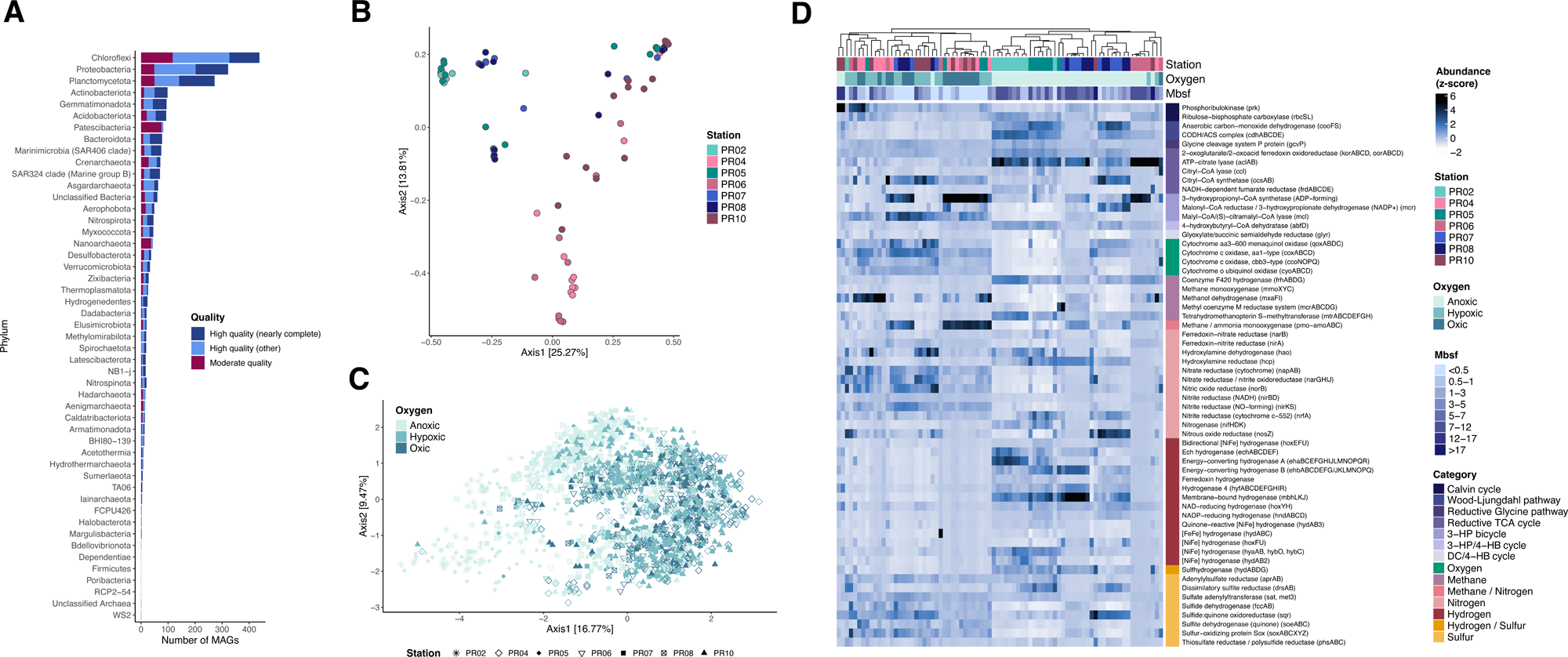
Number of MAGs recovered for different microbial phyla (A), principal coordinate plot of MAG relative abundances converted into Hellinger-transformed Bray-Curtis dissimilarities (B) and differences in functional potential between sedimentary communities (C, D). MAGs are primarily composed of taxa from the phyla Chloroflexi and Proteobacteria (A). Similar to patterns based on ASV abundances, ordination analyses based on MAG abundances (B) indicate structuring of sediment communities into three distinct clusters: 1) shelf and upper slope sediment with little oxygen penetration (PR02, PR05), 2) hadal sediment with little oxygen penetration (PR07, PR08), 3) slope and abyssal-plain sediment with high oxygen penetration (PR04, PR06, PR10). Subseafloor communities also show variation in metabolic potential between sampling stations and oxygen availability in the sediment based on both principal coordinate plots of KEGG module completeness estimated from MAGs (C) and heatmaps of normalized read coverage for key metabolic genes calculated from metagenomes (D). The heatmap was generated based on z-score scaled Euclidean distances of the normalized read counts. A heatmap showing read abundance of metabolic pathways is shown in Fig. S13.

Consistent with the ASV-based analyses, the largest numbers of MAGs were recovered for the phyla Chloroflexi (438) and Proteobacteria (322), followed by Planctomycetota (272) (Fig. 3A), with MAG relative abundances being significantly influenced by station, oxygen availability and sediment depth (Fig. 3B; Table S5; PERMANOVA: p = 0.001, R^2^ = 0.0455–0.2237). Although the sedimentary communities strongly overlapped in metabolic potential based on KEGG module completeness, some functional differences can be observed (Fig. 3C; Table S6, S7; PERMANOVA: p = 0.001, R^2^ = 0.0074–0.0238). For example, MAGs from the bathyal and hadal, mostly anoxic stations show enrichment in characteristic anaerobic processes, such as acetogenesis, Wood-Ljungdahl pathway, methanogenesis (bathyal), dissimilatory nitrate reduction (hadal), and anaerobic benzoate degradation (hadal) (Table S6). By contrast, MAGs at the slope and abyssal plain stations with deep oxygen penetration show overrepresentation of primarily aerobic processes, including thiosulfate oxidation, beta oxidation, cellular respiratory chain, aerobic degradation of phenylacetate and phthalate, and oxygen-dependent biosynthetic pathways (Table S6).

Detailed genomic investigations of select taxonomic groups underscore the functional distinctions between sediment communities at predominantly oxic or anoxic stations (Tables S8-S12; Supplementary Results). For instance, Dehalococcoidia, Caldatribacteriota and Bathyarchaeia, which represented key taxa in anoxic sediment of the Puerto Rico Trench, display potential for versatile anaerobic mixotrophy. Anaerobic members of the Dehalococcoidia, many of which belong to GTDB order Dehalococcoidales, likely grow via oxygen-independent degradation of peptides, amino acids and aromatic compounds (e.g., toluene, benzoate) and, in some cases, autotrophic carbon fixation through the Wood-Ljungdahl pathway (Table S9; Supplementary Results). On the other hand, the almost exclusively hadal Caldatribacteriota (JS1) show capabilities for autotrophic growth via the reductive glycine pathway and heterotrophic metabolism through peptide and sugar fermentation (Table S8; Supplementary Results), with secondary fermentation of metabolic byproducts likely occurring in specialized bacterial microcompartments. Multiple members of this group further demonstrate potential for the biosynthesis of piezolytes, such as trehalose and di-myo-inositol-phosphate, indicating possible adaptation to hydrostatic pressure (28–30). Lastly, Bathyarchaeia, which are predominant in anoxic sediment of the Puerto Rico shelf, appear to utilize the archaeal Wood-Ljungdahl pathway both for autotrophic carbon fixation and assimilation of organic compounds, such as acetate, formaldehyde and methylamines (Table S10; Supplementary Results).

Crenarchaeota and Alphaproteobacteria of the orders Nitrosopumilales and Defluviicoccales, respectively, were characteristic for the oxygenated sediments on the Puerto Rico slope, the mid-trench high and abyssal plain and show potential for aerobic metabolism. Nitrosopumilales are ammonia-oxidizers that use the 3-hydroxypropionate/4-hydroxybutyrate cycle for carbon fixation (Table S11; Supplementary Results), while Defluviicoccales are facultative aerobic heterotrophs that can degrade a broad range of organic substrates, especially nitrogen-containing compounds and possibly halogenated acids (Table S12; Supplementary Results). Members of the Defluviicoccales further show potential for accumulation of polyphosphates, suggesting that they might build carbon and energy reserves for use under unfavorable conditions (Table S12; Supplementary Results).

### Metabolic potential of sediment communities based on metagenomes

Functional analyses based on the unbinned metagenomes broadly agree with the results of the MAG-based analyses, confirming significant metabolic differentiation of microbial communities by station, oxygen availability and sediment depth (Fig. 3D, S13; Table S7; PERMANOVA: p = 0.001, R^2^ = 0.0608–0.1349). Functional differences are especially related to energy, macromolecule, xenobiotics, glycan and secondary metabolism. For example, the coverage of key enzymes for different carbon fixation pathways varies with dissolved oxygen concentration in the sediment, with enzymes for the reductive TCA cycle and Wood-Ljungdahl pathway being more abundant in anoxic sediment and enzymes for the 3-hydroxypropionate bicycle being more prevalent in oxic/hypoxic sediment. Microbial communities in anoxic sediment also show higher abundances of key enzymes for methanogenesis (especially at bathyal depths), nitrous oxide and hydroxylamine reduction (especially at hadal depths), hydrogen metabolism, and dissimilatory sulfite reduction and demonstrate elevated levels of glycan biosynthesis pathways (Fig. 3D, S13). By contrast, microbial communities in oxygenated sediment exhibit higher abundances of key genes for the electron transport chain, nitrate reduction, methane and ammonia oxidation, methylotrophy, and sulfur oxidation through the SOX pathway (Fig. 3D). These communities are further characterized by a prevalence of pathways for degradation of different xenobiotics and biosynthesis of secondary metabolites, especially in the first few decimeters of sediment (Fig.S13).

## Discussion

Our results show that the taxonomic and functional composition of the Puerto Rico Trench deep biosphere is strongly influenced by seafloor topography. Seafloor topography controls where sediment accumulates, in turn impacting the redox chemistry of the porewaters. Consequently, topographic features that accumulate sediment quickly are characterized by anoxic conditions just below the seafloor surface, whereas topographic features that accumulate sediment slowly can remain oxic for many meters below the seafloor surface. These topography-influenced biogeochemical dynamics result in the selection of distinct subseafloor communities in the Puerto Rico Trench. Our results expand prior studies investigating the influence of seafloor topography on hadal microbiomes (13, 15, 21–23), demonstrating that topographic effects are not just restricted to microbial communities living at the sediment surface. Namely, our findings indicate that microbes persisting deep below the seafloor are still impacted by physical constraints of seafloor topography even after a relatively long period of time since deposition.

Although the sedimentary communities are broadly dominated by Chloroflexi, Proteobacteria, Crenarchaeota, Bacteroidota, Caldatribacteriota and Planctomycetota, abundances of these and other taxa vary with sampling station and the associated differences in topography and sediment geochemistry. We identified three main community clusters: 1) bathyal shelf and upper slope sites with low oxygen penetration characterized by high abundances of Chloroflexi (Dehalococcoidia) and prevalence of Bathyarchaeia toward the Puerto Rico shelf (PR02, PR05), 2) hadal sites with low oxygen penetration characterized by high abundances of Dehalococcoidia and predominance of Caldatribacteriota (JS1) in deeper sediment sections (PR07, PR08), and 3) bathyal and abyssal sites with deep oxygen penetration characterized by high frequencies of Alphaproteobacteria, Nitrososphaeria and Nanoarchaeia (PR04, PR06, PR10). This finding supports work from other deep-sea systems that indicate water depth (i.e., hydrostatic pressure) (17, 19, 21) and the availability of dissolved oxygen (31) as fundamental drivers of marine sedimentary microbial communities. The dissolved oxygen concentration in subseafloor sediment depends primarily on the rate of sediment accumulation (32). Abyssal plains and steep trench slopes are characterized by low sedimentation rates that limit burial and preservation of labile organic matter. These characteristics favor lower respiration rates and, thus, deep oxygen penetration into the sediment. Deep oxygen penetration appears to be amplified at the slope site PR04 and the mid-trench high site PR06, where stratigraphic unconformities resulting from periods of erosion and/or non-deposition likely brought older sediment into contact with oxygen-rich bottom water (27). By contrast, regular pulses of sediment influx that characterize continental shelves and the bottom of hadal trenches lead to rapid organic matter burial and thereby promote high subseafloor sedimentary respiration (33). In such environments, oxygen gets depleted within centimeters to decimeters below the seafloor, resulting in mostly anoxic sediment, such as that observed for the shelf, upper slope and hadal sites investigated here.

Despite being both characterized by anoxia, the bathyal shelf and upper slope sites and the hadal trench sites showed distinctions in community composition. This could be related to contrasts in water depth among these sites but might also be caused by differences in sediment geochemistry. For example, the presence of alternative electron acceptors, such as manganese and iron, likely contributes to microbial community structure (13–17, 34), although additional factors, such as continuous, long-term burial and differential mortality of taxa within the subsurface (35–37), might play a role as well.

Changes in taxonomic composition between sedimentary communities are mirrored by changes in metabolic potential across stations, oxygen conditions and subsurface depths. For example, heterotrophic metabolism, aerobic chemolithotrophy and some oxygen insensitive carbon fixation pathways appear to dominate in oxygen-rich conditions prevailing generally at the slope and abyssal plain sites. These sites are characterized by relatively high abundances of ammonia-oxidizing archaea of the order Nitrosopumilales and aerobic heterotrophic Alphaproteobacteria of the order Defluviicoccales. Nitrosopumilales rely on the highly energy-efficient 3-hydroxypropionate/4-hydroxybutyrate cycle for carbon fixation (38), while also being able to use some organic substrates for energy generation (39). Defluviicoccales on the other hand can degrade a variety of organic substances, including many nitrogen-containing compounds and halogenated acids, and demonstrated a feast-or-famine strategy resembling that of polyphosphate accumulating organisms. The capacity for degradation of diverse nitrogenous compounds in Defluviicoccales could suggest that these organisms contribute to ammonia regeneration for the chemotrophic metabolism of Nitrosopumilales, thereby maximizing the use of energy resources. These metabolic characteristics are consistent with the extreme energy and organic carbon limitation of oxygen-rich deep-sea habitats.

By contrast, anaerobic chemolithotrophy, fermentative metabolism, and oxygen sensitive carbon fixation pathways are prevalent under the anoxic subseafloor conditions prevailing generally at the shelf, upper slope and hadal sites. Both uppermost slope and hadal stations are dominated by different groups of Dehalococcoidia, whereas the Puerto Rico shelf area have high frequencies of Bathyarchaeia. Many of these taxa appear to be flexible mixotrophs that can ferment a range of organic carbon compounds but also assimilate carbon dioxide via the Wood-Ljungdahl pathway (40–44), consistent with varying levels of organic carbon and the absence of oxygen in these habitats. In addition, some lineages of Caldatribacteriota typify the deep subsurface sediment at the hadal sites. The occurrence of these taxa coincides with an interval of elevated TOC and TN content and a correlated shift in δ^13^C_org_ values from about -22‰ to -27‰, consistent with slumping and rapid burial of terrigenous slope sediment after a mass wasting event (45). Organic matter enrichment in these sediment sections likely selects for high abundances of Caldatribacteriota over other microbial taxa by fueling their highly efficient fermentative metabolism. Caldatribacteriota are able to conserve additional energy through secondary fermentation in bacterial microcompartments (46, 47), while showing evidence for hydrostatic pressure adaptation (29, 30), thereby likely having a competitive advantage in hadal deep subsurface sediments.

In summary, our data show that seafloor topography influences the composition and metabolic potential of subseafloor microbial communities of the Puerto Rico Trench. First, the subseafloor community in sediment at hadal depths at the bottom of the trench differs from the anaerobic subseafloor community in sediment at bathyal depths, in accordance with the effects of water-depth-related physical properties, such as pressure and temperature, on subseafloor community composition. Second, aerobic communities persist meters beneath the seafloor at lower bathyal and abyssal depths in and near the trench, where sediment accumulation is likely low, leading to deep oxygen penetration into the sediment. This contrasts with the upper bathyal and hadal sites, where sediment accumulation is probably higher and the subseafloor sediment becomes anoxic centimeters to decimeters below the seafloor. These findings advance our understanding of how seafloor topography shapes the composition and functions of microbial communities beneath the seafloor.

## Materials and Methods

### Sample collection

Sediment samples were collected with piston, gravity and multi-corers from seven sites (493– 8385 mbsl) in and around the Puerto Rico Trench during R/V *Neil Armstrong* cruise AR64-02. Cores were sectioned on board the ship immediately after retrieval. Samples for environmental genomics were taken with sterile 60 ml syringes from core-section interiors, taking care to avoid contact with the core liner to reduce contamination. Comparative seawater samples were collected at all sites across the water column with either a CTD or the Deep Autonomous Profiler and filtered shipboard through 0.22 µm Sterivex filters. All samples were frozen immediately at - 80°C after processing.

### Geochemical analyses

Dissolved oxygen, dissolved nitrate, dissolved sulfate and alkalinity, were measured as previously described (48, 49). TOC and δ^13^C_org_ measurements were performed according to ref. (50), while TN and calcium carbonate were measured following ref. (51). As a proxy for active microbial biomass, bulk adenylates were determined according to ref. (52).

### DNA extraction

Genomic DNA from >10 g of sediment per sample was purified using a custom extraction protocol modified after ref. (47) (Supplementary Methods). DNA from seawater samples was extracted using the Qiagen DNeasy PowerWater Sterivex kit (Qiagen, Hilden, Germany) with a vacuum manifold system. Extraction blanks were included in all sediment and seawater sample extractions to assess cross-contamination in the laboratory. All DNA extracts were quantified with the Qubit dsDNA HS assay kit (Thermo Fisher Scientific, Waltham, MA, USA).

### 16S rRNA amplicon sequencing and analysis

Barcoded 16S V4-V5 amplicon libraries were prepared with the 515F/926R primer pairs (53) for 166 sediment samples, 105 seawater samples and 20 negative controls and 2x250 bp paired-end sequenced to an average depth of 583,176 total reads per sample on a NextSeq 2000 platform (Illumina, San Diego, CA, USA) at the Argonne National Laboratory (Lemont, IL, USA) (Table S1).

Raw 16S rRNA amplicon reads were quality-checked with FastQC (https://github.com/s-andrews/FastQC) and then processed with fastp (54) and Trimmomatic (55) to remove polyG tails and sequencing adaptors. Trimmed reads were merged, filtered and denoised into ASVs with Usearch (56) and Vsearch (57). ASV abundances were quantified with the *vsearch -- usearch_global* command using an identity threshold of 99%. ASV sequences were taxonomically classified in Qiime2 (58) with a naïve Bayes classifier weighted for marine sediment and seawater and trained on SILVA 138.1 database sequences trimmed to the V4-V5 primer region. A mid-point rooted phylogenetic tree of the ASV sequences was obtained with FastTree2 (59) in Qiime2 (58). The resulting dataset was filtered for low-abundance ASVs, contamination and read support (Supplementary Methods).

Abundance and ordination plots were constructed with the Phyloseq (60) package in R (61). To assess the influence of environmental factors on microbiome composition we further performed PERMANOVAs, testing marginal effects of each term and multiple distance metrics (Supplementary Methods). Differential abundance of ASVs between seawater and sediment communities and between sediment community clusters identified in PCoAs was determined with DESeq2 (62).

### Metagenomic sequencing and analysis

Low-input metagenomic libraries for 80 select sediment samples were prepared at the University of California Davis and 2x150 bp paired-end sequenced to an average depth of 277,309,556 total reads per sample on a NovaSeqX system (Illumina, San Diego, CA, USA) (Table S1).

Raw reads were trimmed and quality-filtered as described above and then assembled with MegaHit (63) using the *--meta-large* preset option recommended for complex microbiomes.

MAGs were binned from the MegaHit assemblies with MetaBAT2 (64) and MaxBin2 (65). A non-redundant set of best-scoring bins was identified for each sample with DAS Tool (66), quality-checked with the CheckM lineage workflow (67), and taxonomically classified with GTDBtk based on the Genome Taxonomy Database release R226 (68). The GTDB classifications were converted to SILVA 138.1 taxonomy for better comparability with the ASV data. MAGs assigned as *Paenibacillus G validus* (GCF_004000985.1) and *Bradyrhizobium* sp002831585 (GCF_002831585.1) were removed as they likely represented environmental contamination. All other moderate- to high-quality MAGs with completeness >50% and contamination <10% were kept for further analysis.

A dereplicated, species-level set of these MAGs was generated with dRep (69) and used to assess per-sample relative abundances with CoverM (70). To infer variation in metabolic potential among sediment samples, MAGs were functionally annotated in Anvi’o (71) based on the KEGG/KOfam databases. Metabolic capabilities and enrichment were predicted between sediment community clusters with the *anvi-estimate-metabolism* and *anvi-compute-metabolic-enrichment* programs for high-quality MAGs only (>75% completeness). Modules with adjusted p-values < 0.05 were considered enriched. High-quality MAGs from select taxonomic groups were selected for an in-depth analysis of metabolic potential.

For comparison, we also functionally profiled the unbinned metagenomes by predicting protein-coding genes with MetaProdigal (72), excluding contigs belonging to the contaminant *Paenibacillus G validus* and *Bradyrhizobium* sp002831585 MAGs. A non-redundant set of protein-coding genes across metagenomes was generated with the MMseqs2 *easy-linclust* workflow (73), annotated with eggNOG-mapper (74) and filtered for eukaryotic annotations.

Read counts of prokaryotic KO terms were estimated with CoverM (70). Gene abundances in each sample were standardized by dividing the gene-specific read counts by the average read count of 35 single-copy COGs defined in ref. (75). We then assigned genes to KEGG metabolic pathways and calculated the average normalized read count for each pathway. Additionally, we estimated average abundances of key metabolic genes for carbon, nitrogen, sulfur, hydrogen, oxygen, and methane metabolism. Genes of interest were selected based on previous studies and functional relevance (75–77). Heatmaps and principal coordinate plots of microbiome functional potential were created with the ComplexHeatmap (78) and ggplot2 (79) packages in R (61).

## Acknowledgements

This research would not have been possible without the dedicated effort of the captain and crew of R/V *Neil Armstrong* expedition AR64-02. We thank Paul Walczak, Christopher Fanshier, Al Gagnon, and Chris Griner for assembling and operating the many coring systems used by this project for sampling the seafloor. We also thank David Casagrande and Kotachi Liu for operating the URI Deep Autonomous Profiler used to take water-column samples for this study. We gratefully acknowledge the Argonne National Laboratory for preparing and sequencing the 16S rRNA amplicon libraries and the University of California Davis Genome Center for preparing and sequencing the metagenomic libraries. This work was primarily funded by the National Science Foundation grants OCE-2049515 to S.D., R.P., and Y.Z. and OCE-2048927 to D.H.B. A portion of this work was funded by DFG project OR417-7-1 to W.D.O. D’Hondt is a CIFAR Fellow in the Earth 4D: Subsurface Science and Exploration Program.

## Data Availability

Raw 16S rRNA amplicon reads, raw metagenomic reads and metagenome-assembled genomes have been deposited into the National Center for Biotechnology Information under BioProject PRJNA1334354. Bioinformatic code used to conduct analyses is available on GitHub under repository https://github.com/cbreusing/PRT_microbiome.

## Author Contributions

C.B. performed molecular laboratory work, analyzed the sequence datasets and wrote the original draft. M.D., A.M.P., W.D.O. and D.H.B. collected sediment samples and discussed results during team meetings. W.D.O. developed the study’s narrative structure, advised on DNA extraction protocols and led adenylate, TN and CaCO_3_ measurements. D.C.S. supported collection of seawater samples and associated metadata. R.P. created bathymetric maps of the Puerto Rico Trench. I.A., E.A.S., A.J.S., and T.W. took porewater samples for post-cruise chemical analyses. D.G., and L.M.W. performed porewater and sediment chemistry analyses, respectively. S.R.S, B.S. and J.J.S.-C. helped with sample collection and processing on the cruise. D.H.B. and S.D. designed the study, interpreted findings during group meetings and, together with Y.Z. and R.P, acquired grant funding. All authors contributed to manuscript writing.

## Classification

Biological Sciences, Microbiology

