## Supplementary Information for "Seafloor topography structures subseafloor microbial communities in the Puerto Rico Trench"

**This PDF file includes:**

Supplementary Methods
Supplementary Results
Supplementary Discussion
Supplementary References
Figures S1 to S13

Supplementary Tables S1 to S12 are provided in a separate Excel file.

**Supplementary Methods**

*DNA extraction protocol for sediments and soils*

1. Prepare C1 extraction buffer. Heat to 65°C to dissolve phosphate crystals. Sterile filter and UV treat (20 min) C1 extraction buffer. Store buffer at room temperature.

   C1 extraction buffer recipe (for 250 mL volume):

   - 190 mL 1 M Disodium hydrogen phosphate solution
   - 37.5 mL 100% Ethanol
   - 20 mL Solution C1 (Qiagen)
   - 1.5 mL 10% SDS

2. UV-treat all non-enzymatic buffers, reagents and other materials before starting with the extractions. Sterilize laminar flow hood with DNA Away solution. Turn on air purifier and static eliminator. Sterile flame subsampling tools and clean with DNA Away solution between all samples.
3. Let samples thaw for about 30 minutes until sediment is soft enough to be subsampled. Cut the first ~1 cm of sediment off, then subsample the core of the sample.
4. Add sample to Lysing Matrix E tube (placed on balance within the laminar flow hood) until desired weight is reached. Add at least 1-1.5 mL pre-heated (65°C) C1 extraction buffer per g of sample. Leave at least 1-2 cm of head space between the top of solution and the top of tube.
5. Homogenize for 80 seconds at a speed of 3.1 m/s on the Omni Bead Ruptor 24.
6. Heat at 99°C for 2 minutes.
7. Freeze at -70°C overnight.
8. Thaw. Make sure phosphate crystals go back into solution by heating at 65°C.
9. Heat at 99°C for 2 minutes.
10. Homogenize for 80 seconds at a speed of 3.1 m/s on the Omni Bead Ruptor 24.
11. Centrifuge at top speed for 5 minutes for 1 mL tubes, or 15 minutes for 50 mL tubes.
12. Concentrate supernatant down to 200-400 µL in 30 kDa MWCO Amicon filters. Heat tubes briefly at 65°C to avoid formation of phosphate crystals during centrifugation. Spin filters in

two 15-minute intervals at 4000 g and 35°C, checking for phosphate crystals in between runs and heating at 65°C for a few minutes if necessary. Then centrifuge for another 10 minutes.

13. Clean up the concentrate with the Omega Bio-Tek EZNA Soil DNA kit (starting at step 7 adjusting volumes of P2 buffer and cHTR reagent). Perform enough inhibitor removal steps so that the supernatant is reasonably clear. Elute in 100 µL.
14. Perform a secondary DNA cleanup with the Qiagen DNeasy PowerClean Pro kit, eluting in 50 µL pre-warmed (70°C) elution buffer. Repeat the elution step using the filtrate of the previous step for elution.
15. Measure on Qubit / Nanodrop and store at -20°C to -80°C.

*Required materials:*

- C1 solution (Qiagen, Cat. No. 12888-100-1)
- 1 M Disodium hydrogen phosphate solution, sterile filtered (Tribioscience, Cat. No. TBS5046)
- 100% Ethanol, Molecular Biology Grade (Fisher Scientific, Cat. No. BP28184AC22)
- 10% Sodium Dodecyl Sulfate Solution, Molecular Biology Grade (Fisher Scientific, Cat. No. BP2436200)
- Thermo Scientific™ Nalgene™ Rapid-Flow™ Sterile Disposable Filter Units (Fisher Scientific, Cat. No. 09-741-04)
- MP Biomedicals™ Lysing Matrix E 50 mL tubes (Fisher Scientific, Cat. No. 11-695-4050). If these tubes are backordered order bulk beads and tubes to prepare lysing matrix:
    - 50 mL impact resistant Falkon tubes (Omni International, Cat. No. 19-6650)
    - Lysing Matrix B beads (MP Biomedicals, Cat. No. 116540425-CF), add 11.25 g per 50 mL tube
    - Lysing Matrix D beads (MP Biomedicals, Cat. No. 116540434), add 18.25 g per 50 mL tube
    - 4 mm glass beads (MP Biomedicals, Cat. No. 112521011), add 25 beads per 50 mL tube
- 30 kDa MWCO MilliporeSigma™ Amicon™ Ultra-15 Centrifugal Filter Units (Fisher Scientific, Cat. No. UFC903008)
- Thermo Scientific™ DNA AWAY™ Surface Decontaminant (Fisher Scientific, Cat. No. 21-236-28)
- EZNA Soil DNA kit (Omega Bio-Tek, Cat. No. D5625-02)
- cHTR reagent (Omega Bio-Tek, Cat. No. cHTR-50)
- Qiagen DNeasy PowerClean Pro Cleanup kit (Qiagen, Cat. No. 12997-50)
- Qubit dsDNA HS Assay Kit (Fisher Scientific, Cat. No. Q32851)
- For challenging samples:
    - 0.1 mm G2 Enhancer Beads (Ampliqon, Cat. No. A420100)
    - 1.4 mm G2 Enhancer Beads (Ampliqon, Cat. No. A421400)

*ASV filtering and decontamination*

Singletons and ASVs with low prevalence and abundance (i.e., less than 10 total reads across less than 5% of samples) were removed from the dataset in R (1) with the PhyloSeq (2) and metagMisc (https://github.com/vmikk/metagMisc) packages. In addition, we excluded one ASV that was identified as a fungal laboratory contaminant (*Komagataella phaffii*) stemming from one of the polymerases used for library preparation. To account for potential cross-contamination in the laboratory, ASV counts in the filtered dataset were corrected by subtracting counts in the corresponding extraction blanks. Samples with less than 50,000 remaining reads were removed from the final dataset.

*Choice of distance metrics for PERMANOVAs and principal coordinate analyses*

We used various distance metrics, including Hellinger-transformed Bray-Curtis dissimilarities and weighted UniFrac distances as well as rarefied Jaccard distances and unweighted UniFrac distances. As equivalent results were obtained for all metrics, but patterns became most apparent using unweighted UniFrac distances, we only present outcomes based on this metric in the manuscript. We provide results for the other metrics in the supplement.

### Supplementary Results

*Sediment chemistry*

The calcium carbonate concentration data are broadly consistent with the three bathyal sites being above the carbonate compensation depth (CCD) and the abyssal and hadal sites being below the CCD. Calcium carbonate content is consistently high throughout the sediment interval at the bathyal sites (PR02, PR04, PR05), with values generally exceeding 50 wt% (Fig. S5). By contrast, it quickly decreases with sediment depth at the mid-trench high (PR06) and the abyssal plain (PR10), approaching zero a couple of decimeters below the seafloor. The $CaCO_3$ profiles of the hadal stations are more complex, with values shifting from near zero at the seafloor to about 50 wt% at ca. 0.2–0.3 mbsf and returning to near zero at ca. 3.5–5 mbsf (Fig. S3).

At all sites, total organic carbon (TOC) content is less than 1 wt%. TOC is overall higher at the bathyal station on the shelf (PR02) and the hadal stations at the trench bottom (PR07, PR08) than at the stations on the slope (PR04, PR05), the mid-trench high (PR06) and the abyssal plain (PR10) (Fig. S6). TOC profiles for the two hadal stations are broadly similar to the profiles at PR02 for the first 5–6 mbsf but reach much higher values for the remaining few meters of the cored sequence (Fig. S1). Organic carbon content is lowest at slope station PR04, with values consistently below 0.25 wt%. The TOC profiles are broadly similar at the upper slope station (PR05), the mid-trench high (PR06), and the abyssal plain station, PR10 (Fig. S6).

The $\delta^{13}C$ values of total organic carbon ($\delta^{13}C_{org}$) at all sites are broadly compatible with production by marine phytoplankton (3). At the bathyal sites closest to shore (PR02, PR05), $\delta^{13}C_{org}$ values are generally heavier than -24‰, with respective median values of -20.85‰ and -22.14‰ (Fig. S7). The $\delta^{13}C_{org}$ signature at the deeper bathyal station PR04 is relatively variable but lighter than at PR02 and PR05, with a median value of -24.6‰. At the two abyssal stations (PR06, PR10), $\delta^{13}C_{org}$ data are even lighter than at the bathyal sites, with a median value of -26‰, (Fig. S7). At the two hadal stations (PR07, PR08), the median $\delta^{13}C_{org}$ value for the first 5–6 mbsf (overlapping with the $CaCO_3$-rich interval) is -22‰, similar to the values at bathyal site PR05. At greater depths below seafloor, however, $\delta^{13}C_{org}$ values are notably lighter, with a median value of about -27‰ (Fig. S7). The relatively light $\delta^{13}C_{org}$ values at the abyssal sites and the lower portion of the cored sediment sequence at the hadal sites are consistent with derivation of the total organic matter from either terrigenous C3 plants (3) or marine phytoplankton biomass synthesized relatively deep in the euphotic zone, as might be expected where sea surface nutrient concentrations are low (4).

Total nitrogen (TN) content declines steeply within decimeters below the seafloor at four of the seven sites (PR04, PR07, PR08, PR10) (Fig. S8). At greater sediment depths at the slope station (PR04) and the abyssal plain station (PR10), TN content is relatively constant. At the two hadal stations (PR07, PR08), TN profiles resemble the TOC profiles, with TN values relatively low for the first few meters below the near-seafloor interval of steep decline and then increasing to higher values beginning at 4–5 mbsf. TN content declines more slowly with depth at the shelf site PR02 and is relatively low, with no significant net change, at upper slope site PR05 (Fig. S8). In contrast to the other stations, TN concentrations at PR06 on the mid-trench high, increase with sediment depth, from the seafloor to the last measurement point at ca. 2.5 mbsf (Fig. S8).

*Differences in KEGG module representation related to sampling station*

MAGs from the bathyal and hadal, mostly anoxic stations show enrichment in acetogenesis, formaldehyde assimilation via the ribulose monophosphate pathway, and the Wood-Ljungdahl pathway (Table S6). MAGs from the shelf and upper slope further exhibit high representation of methanogenesis, incomplete reductive TCA cycle, pentose (bis)phosphate pathway, acetyl-CoA pathway, semi-phosphorylative Entner-Doudoroff pathway and archaeal biosynthesis of C5 and C10-C20 isoprenoids, while MAGs from hadal stations are enriched in dissimilatory nitrate reduction, benzoate, lysine and arginine degradation and biosynthesis of different amino acids (Table S6). By contrast, MAGs at the slope and abyssal plain stations with deep oxygen penetration show high representation of nitrate assimilation, thiosulfate oxidation, assimilatory sulfate reduction, Calvin and reductive TCA cycles, respiratory complexes, heterotrophic metabolism (e.g., degradation of different amino acids, fatty acids, purines, phenylacetate and phthalate), as well as a variety of biosynthetic pathways, some of which functions under strictly or primarily aerobic conditions (e.g., ubiquinone biosynthesis, NAD biosynthesis from tryptophan, ketogenesis, glycine-dependent thiamine biosynthesis) (Table S6).

*Detailed description of metabolic capacities identified in key microbial community members*

**Caldatribacteriota (JS1).** Members of the JS1 lineage are particularly frequent in deep sediment layers at the bottom of the Puerto Rico Trench. These bacteria appear to be anaerobic mixotrophs that have been shown to grow both through autotrophic carbon fixation and fermentation of sugars and n-alkanes (5–8). In contrast to JS1 taxa from oil reservoirs and abyssal sediment (6, 8), most MAGs investigated here do not encode genes for the carbon monoxide dehydrogenase (CODH)/acetyl-CoA synthase (ACS) complex, indicating that they may not primarily rely on the Wood-Ljungdahl pathway for carbon fixation (Table S8). Instead, like other Caldatribacteriota lineages some of the JS1 MAGs from the present study seem to be able to assimilate carbon through the reductive glycine pathway, for which key enzymes were detected (fdh, fhs, folD, gcvT, gcvH, gcvPAB, glyA, sdaAB). The presence of a monomeric carbon monoxide dehydrogenase (cooS) suggests that the JS1 MAGs might have the potential to use CO as a sole source of carbon and energy. Characteristic genes encoding fumarate addition enzymes critical for short-chain n-alkane degradation were not found in the JS1 MAGs. However, transporters and enzymes for the acquisition and utilization of carbohydrates, peptides and amino acids are present, suggesting potential for sugar and peptide fermentation (Table S8). Energy from fermentative reactions might be conserved through a membrane-bound hydrogenase complex (mbhJKL). As other Caldatribacteriota lineages, the JS1 MAGs from this study contain genes for bacterial microcompartments (eutN, eutM, eutE, DERA), which have been hypothesized to enable secondary fermentation of metabolic byproducts for increased ATP yields under extreme energy limitation (8). Most of the JS1 MAGs further encode genes involved in the synthesis of trehalose (treT) and di-myo-inositol-phosphate (dipps, ino1), two solutes that might be important for adaptation to environmental stress, including hydrostatic pressure (5, 9).

**Chloroflexi (Dehalococcoidia).** Members of the Dehalococcoidia dominate at both bathyal and hadal stations with low oxygen penetration. Taxa in this group have been found to play an important role in the degradation of recalcitrant organic matter, in particular halogenated and aromatic compounds (10, 11). Analyses of the Puerto Rico Trench Dehalococcoidia MAGs provide evidence for a versatile metabolism, with distinctions between taxa that occur in oxygenated versus oxygen-limited sediment (Table S9). MAGs from the slope and abyssal plain stations and the oxic hadal surface sediment, many of which belong to GTDB orders Australimonadales, Lucifugimonadales, DSTF01 and SM23-28-2, appear to be aerobic heterotrophs encoding key genes for oxidative phosphorylation (e.g., coxABCD) and the oxygen-dependent degradation of organosulfur compounds (ssuD), phthalate (pht3, pht4, pht5) and halocarbons (dhaA, dehH). By contrast, MAGs from deeper hadal sediment sections and the shelf and upper slope stations, many of which belong to GTDB order Dehalococcoidales, demonstrate potential for anaerobic respiration, especially based on CO (cooFS) and $H_2$ (hoxFU), as well as the oxygen-independent degradation of aromatic compounds, such as toluene (bbsB, bbsEF, bbsH) and benzoate (e.g., bcrABCD). In addition, some of these MAGs contain genes for reductive dehalogenation (e.g., pceA), indicating capacity for organohalide respiration using hydrogen as electron donor. Potential for consumption of peptides and amino acids (e.g., pepA, pepQ, ilvE) is present across all Dehalococcoidia MAGs. Interestingly, several of the Dehalococcoidia MAGs from anaerobic sediment encode genes for the bacterial-type Wood-Ljungdahl pathway, while MAGs from oxic sediment encode genes for the reductive glycine pathway, which suggests that some of the recovered taxa may have the ability for autotrophic growth.

**Crenarchaeota (Bathyarchaeia).** Members of the Bathyarchaeia are most abundant at the shallowest station on the Puerto Rico shelf and have previously been identified as opportunistic, anaerobic mixotrophs that can rely on both chemosynthetic carbon fixation and methylotrophic/acetogenic fermentation for growth (12–14). Genome investigations of the corresponding MAGs indicate that members of this group can assimilate carbon dioxide mainly through the archaeal-type Wood-Ljungdahl pathway (Table S10), which is likely powered by an electron-bifurcating protein complex consisting of heterodisulfide reductase (hdr) and F420 hydogenase (mvh). In this complex, F420 hydrogenase oxidizes molecular hydrogen to provide reducing equivalents to heterodisulfide reductase, which then splits these electrons between high-potential electron acceptors and low-potential ferredoxin to drive the thermodynamically unfavorable reactions of the Wood-Ljungdahl pathway. The potential for a heterotrophic lifestyle is present through use of acetate and methylated carbon compounds, such as formaldehyde and methylamines, which can be assimilated or respired through the Wood-Ljungdahl pathway via acetyl-CoA synthetase (acs), 5,6,7,8-tetrahydromethanopterin hydro-lyase (fae) and trimethylamine---corrinoid protein Co-methyltransferase (mttB).

**Crenarchaeota (Nitrososphaeria, Nitrosopumilales).** Members of the Nitrosopumilales were mainly observed in deep-sea oxic/hypoxic sediments. Taxa in this group are known aerobic ammonia-oxidizing archaea that assimilate carbon dioxide through the 3-hydroxypropionate/4-hydroxybutyrate cycle (15), as evidenced by the presence of key enzymes for nitrification (e.g., methane / ammonia monooxygenase [pmo-amoABC], nitrite reductase [nirK]), and carbon fixation (e.g., 4-hydroxybutyryl-CoA dehydratase [abfD]) in their genomes (Table S11). Some Nitrosopumilales MAGs further encode genes for transport and degradation of urea (ureABC, urtA), cyanate (cynS), peptides and amino acids (e.g., ABC.PE, pepA, pepF, pepQ), suggesting they may be able to use these nitrogen compounds for ammonia regeneration in ammonia-limited environments. In addition, the presence of formate dehydrogenase (fdh) and pyrroloquinoline quinone dependent aldose sugar dehydrogenase (yliI) indicates that some Nitrosopumilales may have the potential to oxidize formate and sugars for energy generation. Surprisingly, none of the recovered Nitrosopumilales MAGs encode genes for glutamate dehydrogenase (gdh), which is supposed to be part of the core metabolism of ammonia-oxidizing archaea (15). However, all MAGs contain genes for glutamine synthetase (glnA) and other key enzymes for amino acid biosynthesis (e.g., aspB, asnB). This could either indicate that gdh genes were missed during binning or annotation or that the nitrogen assimilation strategy in the Puerto Rico Trench Nitrosopumilales is divergent from other ammonia-oxidizing archaea.

**Proteobacteria (Alphaproteobacteria, Defluviicoccales).** Members of the Defluviicoccales are most frequent at the slope and abyssal plain stations with deep oxygen penetration. Taxa within this group have been found to be versatile aerobic heterotrophs that can use a variety of organic compounds for growth (16, 17). Genomic investigations of the corresponding MAGs indicate that members of the Defluviicoccales have the potential to take up and degrade a broad range of substrates, though in particular amino acids (e.g., livFGHKM, ABC.PA, bztABCD, ilvE), peptides (e.g., oppF, ABC.PE, dppDF, pepQ, amiE), polyamines (e.g, ABC.SP, potABCD, spuC), urea (urtABCDE, ureABC) and other nitrogen-containing compounds (e.g., ptsN, nthAB) (Table S12). Similar to other representatives of Defluviicoccales (17), the MAGs studied here lack genes for the glyoxylate shunt but encode genes for the ethylmalonyl-CoA pathway (e.g., croR, ccr, epi, mcd) that enables growth on acetyl-CoA generating compounds. In addition, detection of membrane-bound formate dehydrogenase (fdh) indicates that some members may use formate as electron donor for energy generation. The widespread presence of cytochrome c oxidase (coxABCD) suggests that respiration is predominantly aerobic, though encoding of enzymes for dissimilatory nitrate and sulfate reduction (e.g., narGHIJ, sat, aprAB, dsrAB) in some Defluviicoccales genomes imply the possible use of nitrate or sulfate as alternative electron acceptors under oxygen-limited conditions. In contrast to the currently available Defluviicoccales genomes from wastewater, which are known glycogen-accumulating organisms (17), the MAGs recovered in this study do not appear to encode characteristic enzymes for glycogen synthesis (glgABC). Instead, a few members encode typical enzymes for the accumulation of polyphosphates (e.g., polyphosphate kinase [ppk], low affinity phosphate transporter [pit]) in addition to key enzymes of the methylmalonyl-CoA pathway (pccAB, epi, mut) and polyhydroxyalkanoate (PHA) synthesis (phaC), indicating they may store polyphosphates and PHAs as sources of energy and carbon, respectively. The majority of Defluviicoccales further encode genes for 2-haloacid and haloacetate dehalogenases, implying potential for biodegradation of organohalides.

**Supplementary Discussion**

Although deep-sea sediment shares microbial taxa with the overlying water column, the two environments contain different microbial communities (18). Compared to sedimentary microbiomes, seawater communities have higher abundances of Proteobacteria, Marinimicrobia and SAR324 clade bacteria (among others) and were enriched in 3,670 microbial taxa across 33 phyla. Microbes inhabiting seawater experience different environmental pressures than those in sediment, which likely explains contrasts in community composition between these habitat types (18, 19). While water depth had a similar effect on seawater community structure as in sediment, no pattern by sampling station was observed. This finding is consistent with other studies from the North Atlantic, Arctic and South Pacific Ocean showing that pelagic microbial assemblages are vertically but not horizontally stratified due to environmental selection across water masses that differ in physico-chemical characteristics, such as temperature, dissolved organic matter concentration and oxygen availability (20–24). This stratification is further intensified by lateral advection favoring dispersal within the same water mass (24). Consequently, it is possible that the depth-related partitioning of seawater communities enhances compositional differences among sedimentary microbiomes by limiting the pool of taxa that is available for recruitment in sediment of a particular depth zone.

### Supplementary Figures

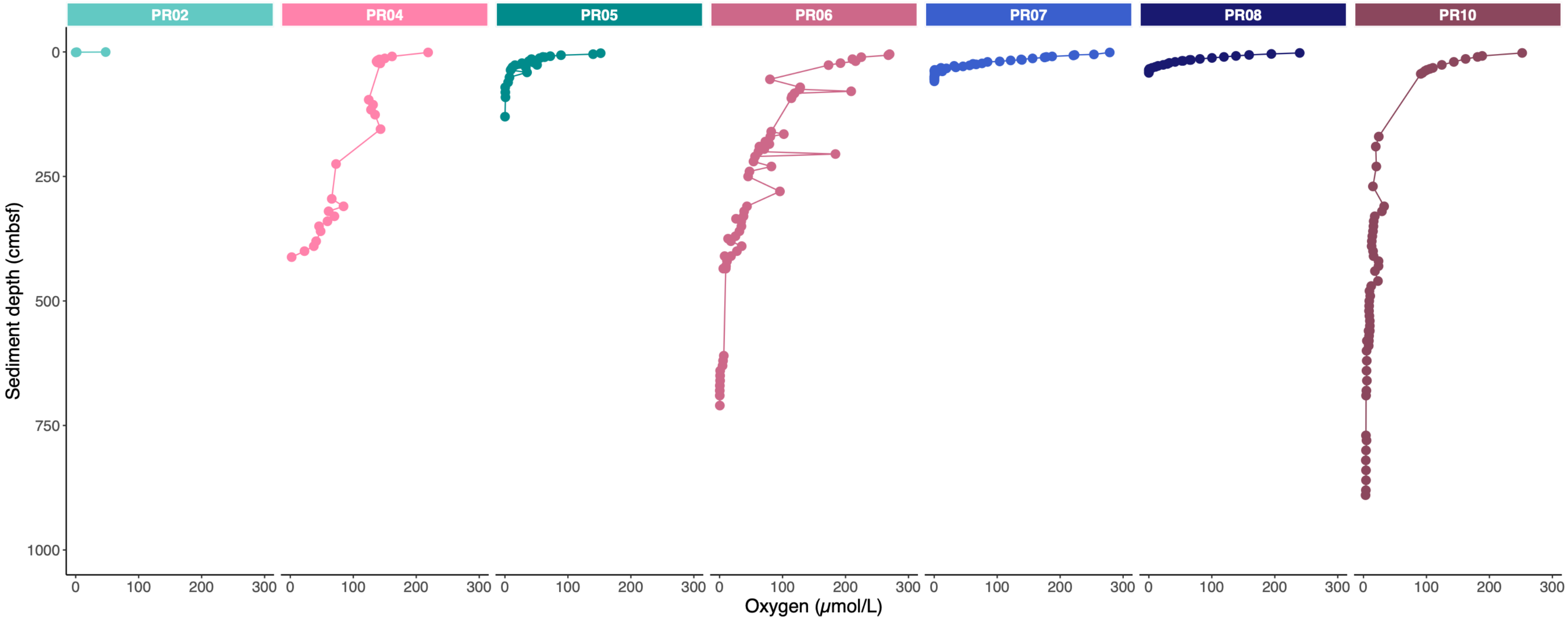


**Fig. S1** Dissolved oxygen profiles.

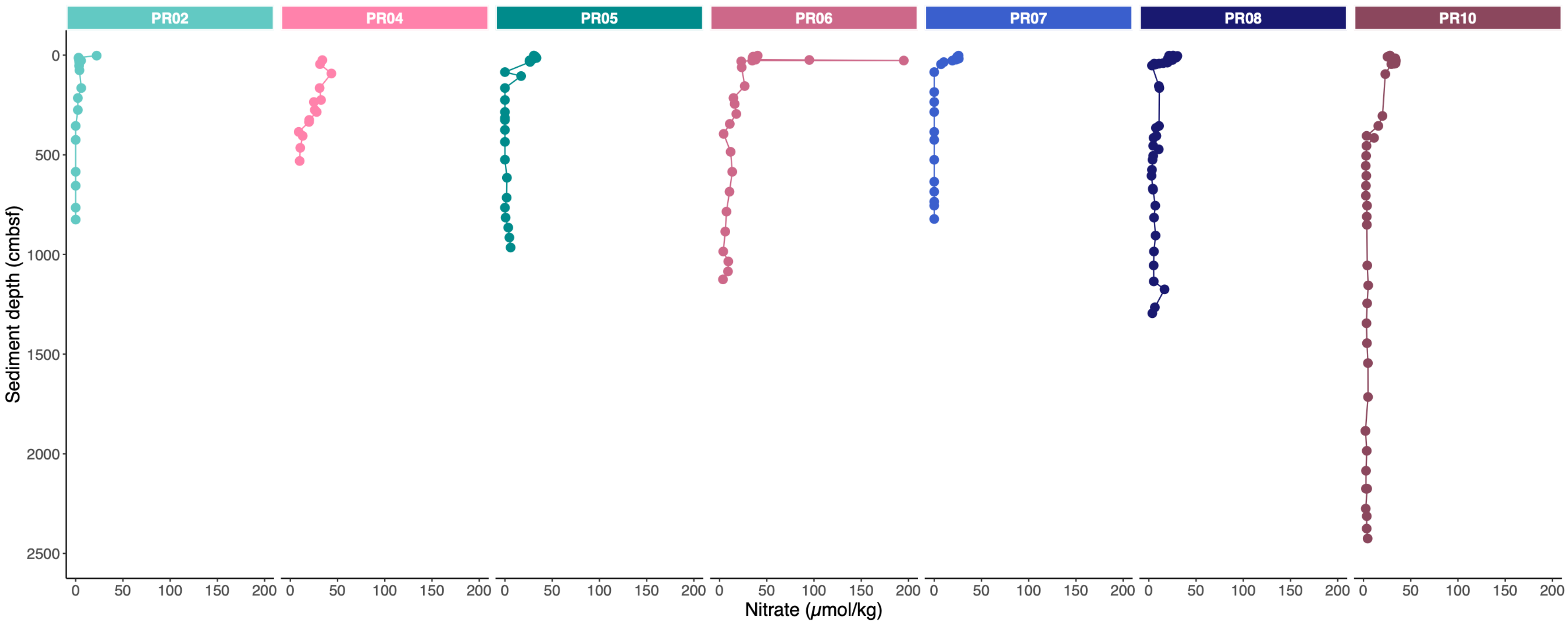


**Fig. S2** Dissolved nitrate profiles.

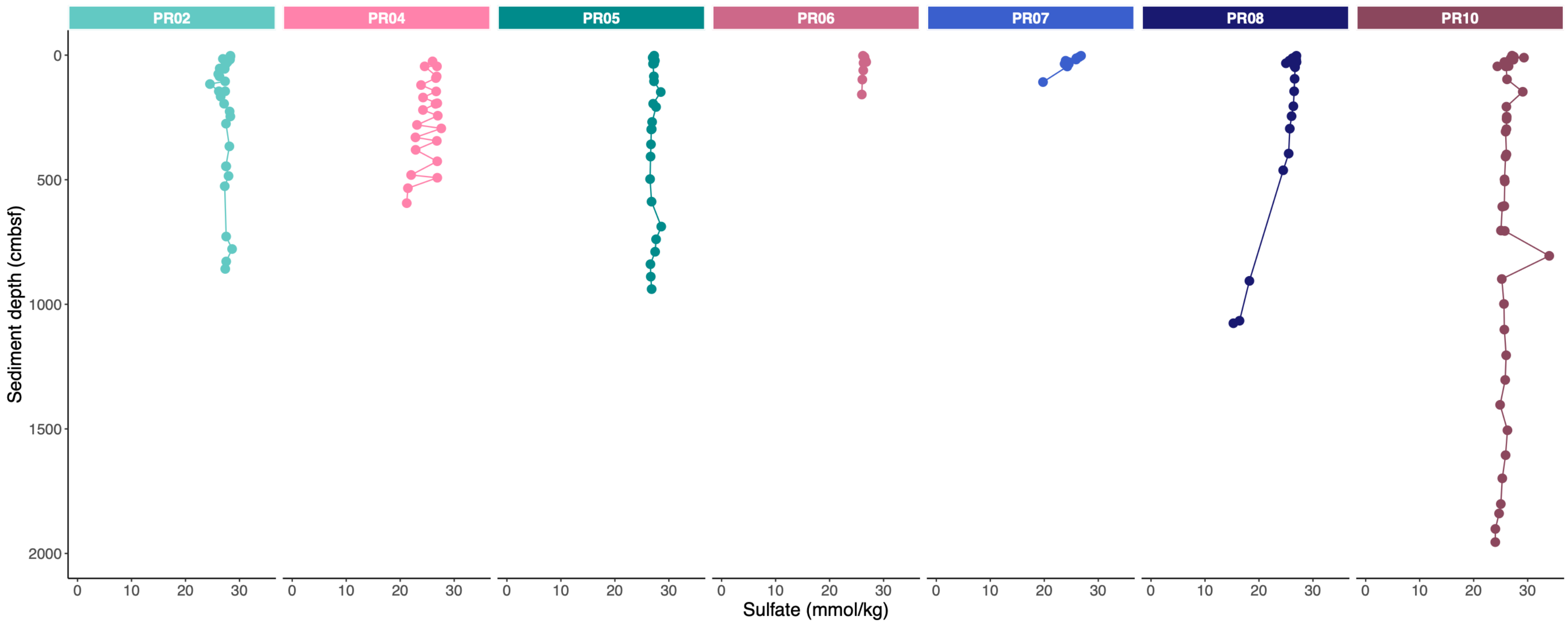


**Fig. S3** Dissolved sulfate profiles.

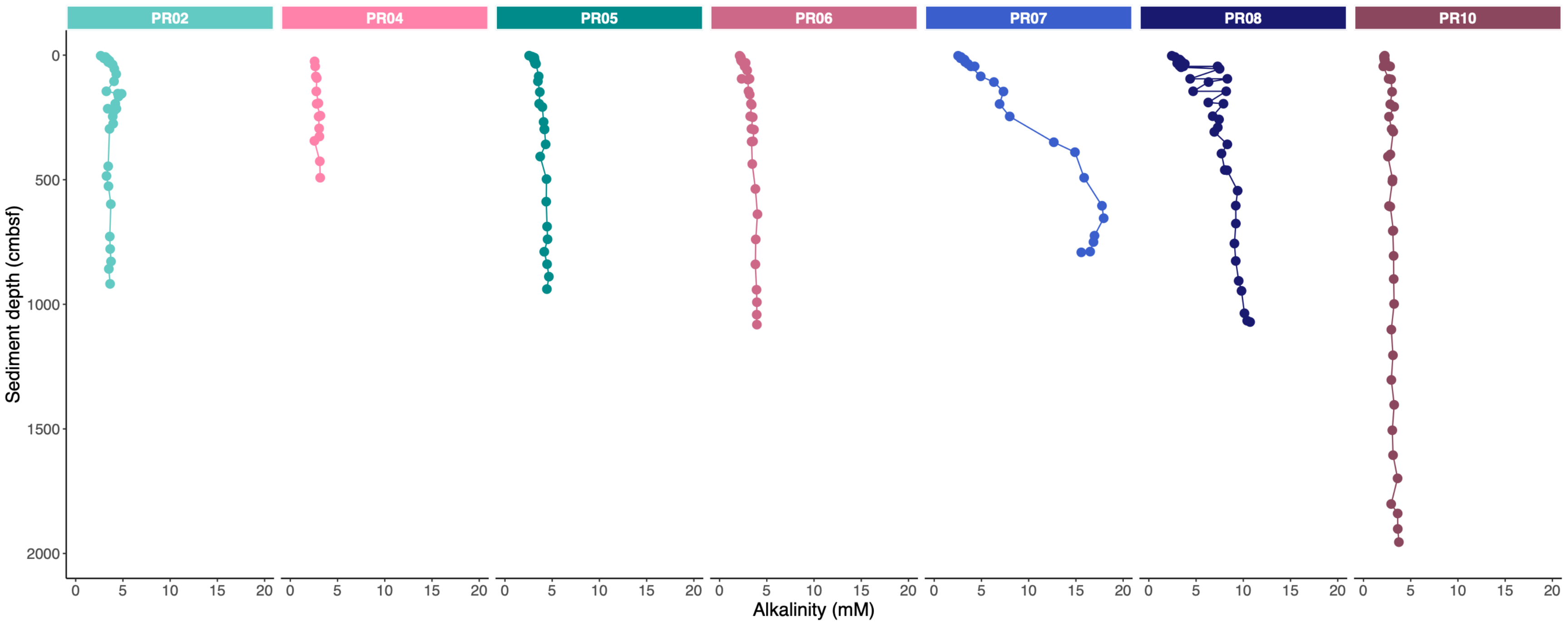


**Fig. S4** Dissolved alkalinity profiles.

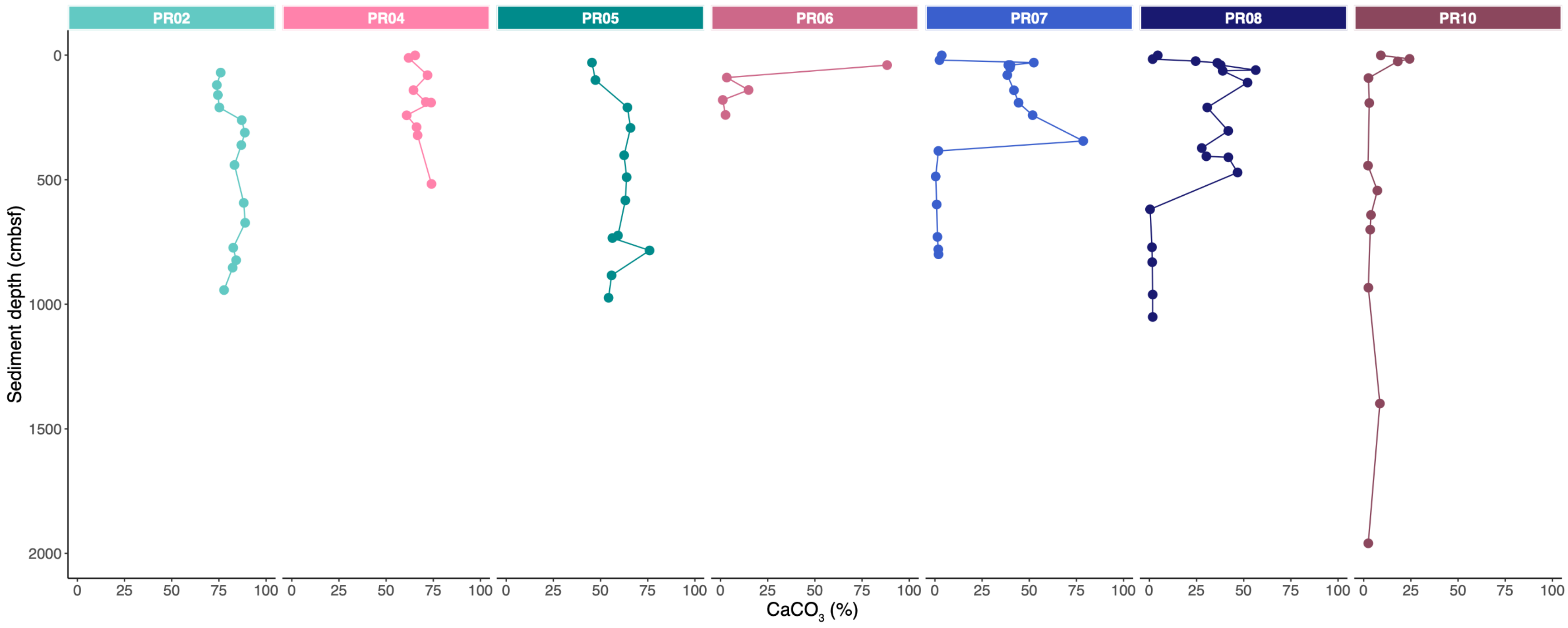


**Fig. S5** Profiles of sedimentary calcium carbonate content.

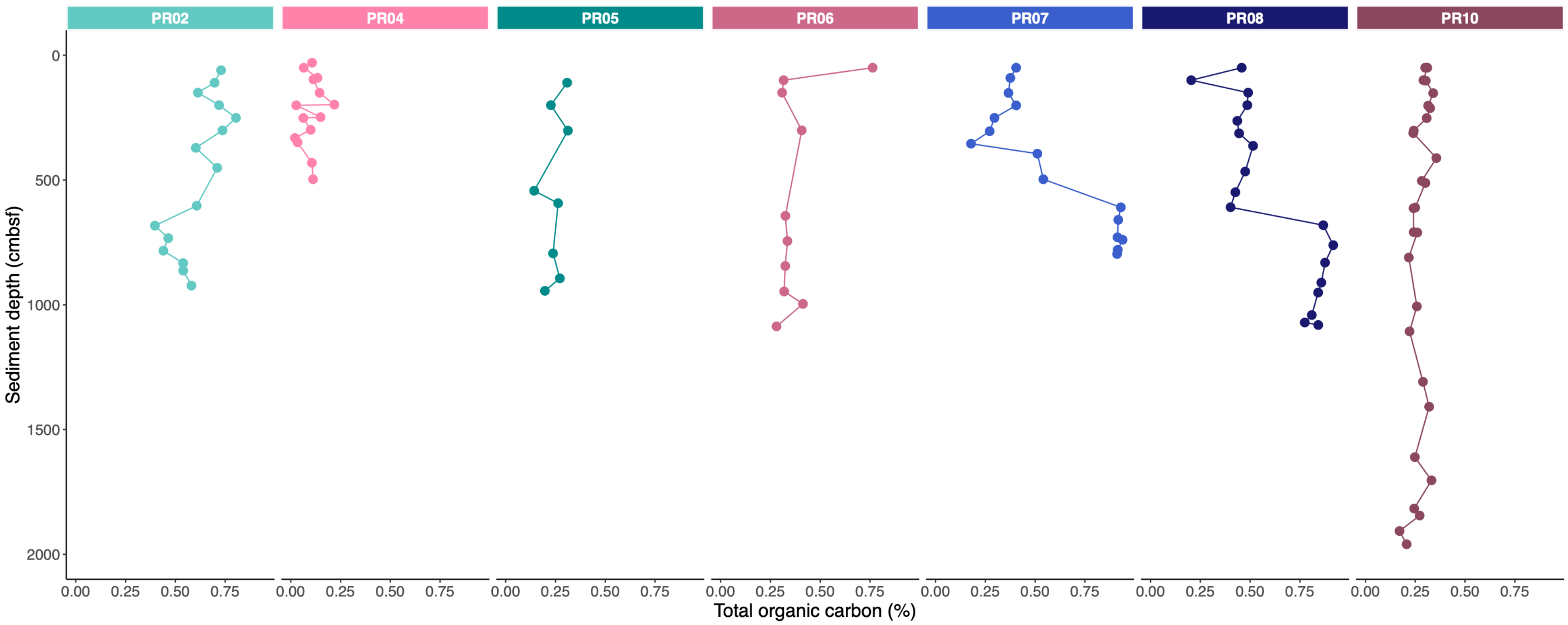


**Fig. S6** Profiles of sedimentary total organic carbon.

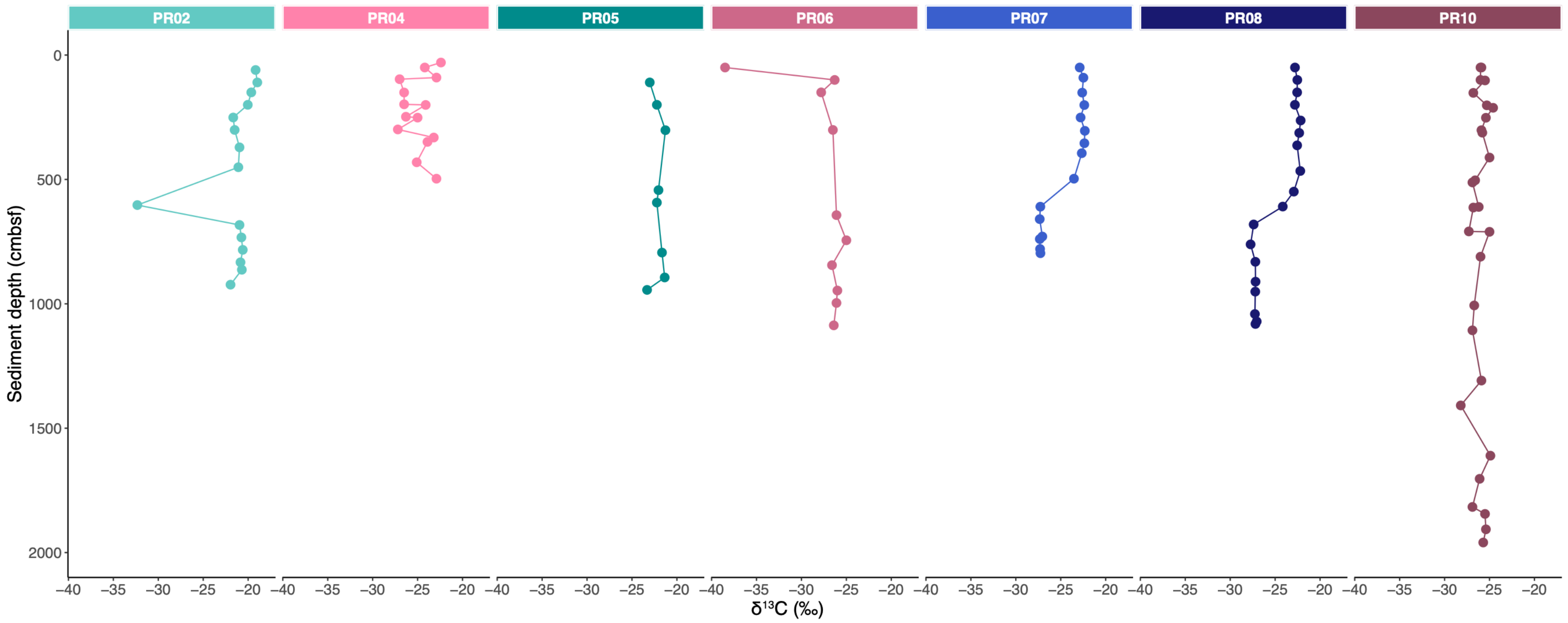


**Fig. S7** Sedimentary profiles of bulk-organic-matter $\delta^{13}C$ values.

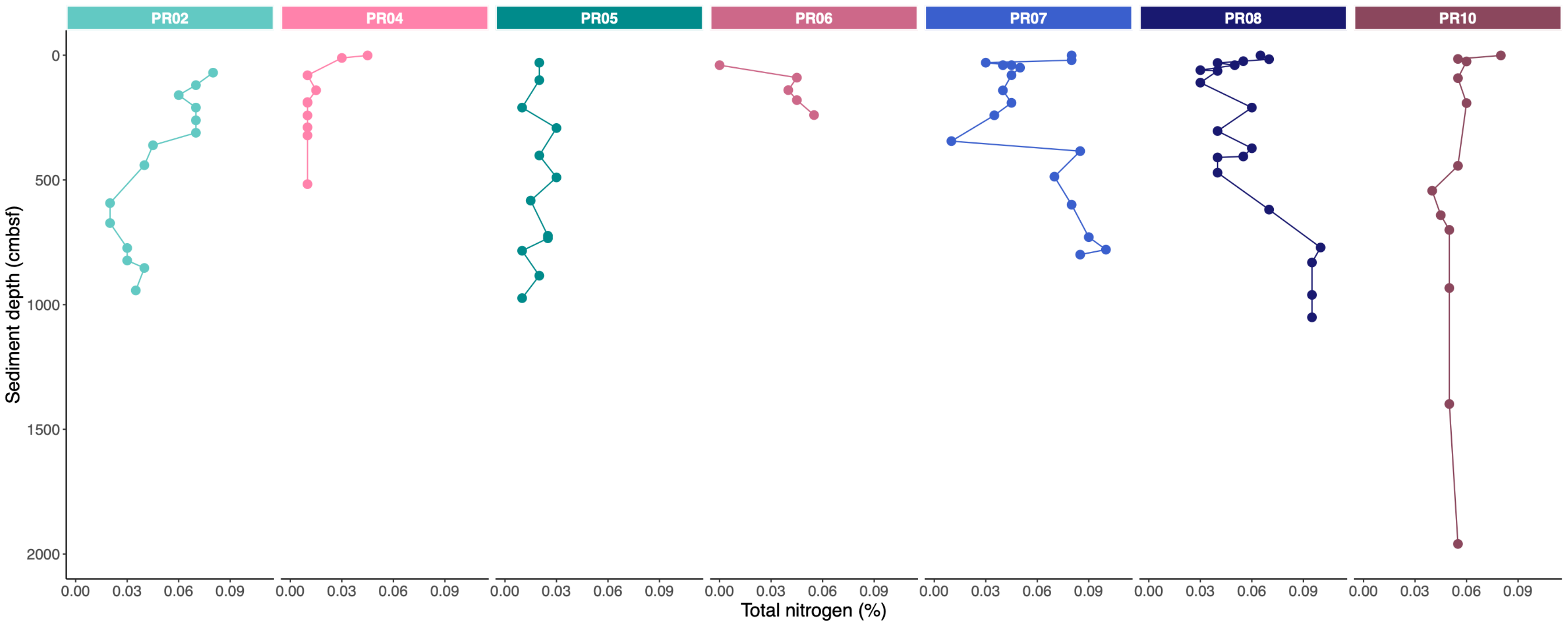


**Fig. S8** Profiles of sedimentary total nitrogen.

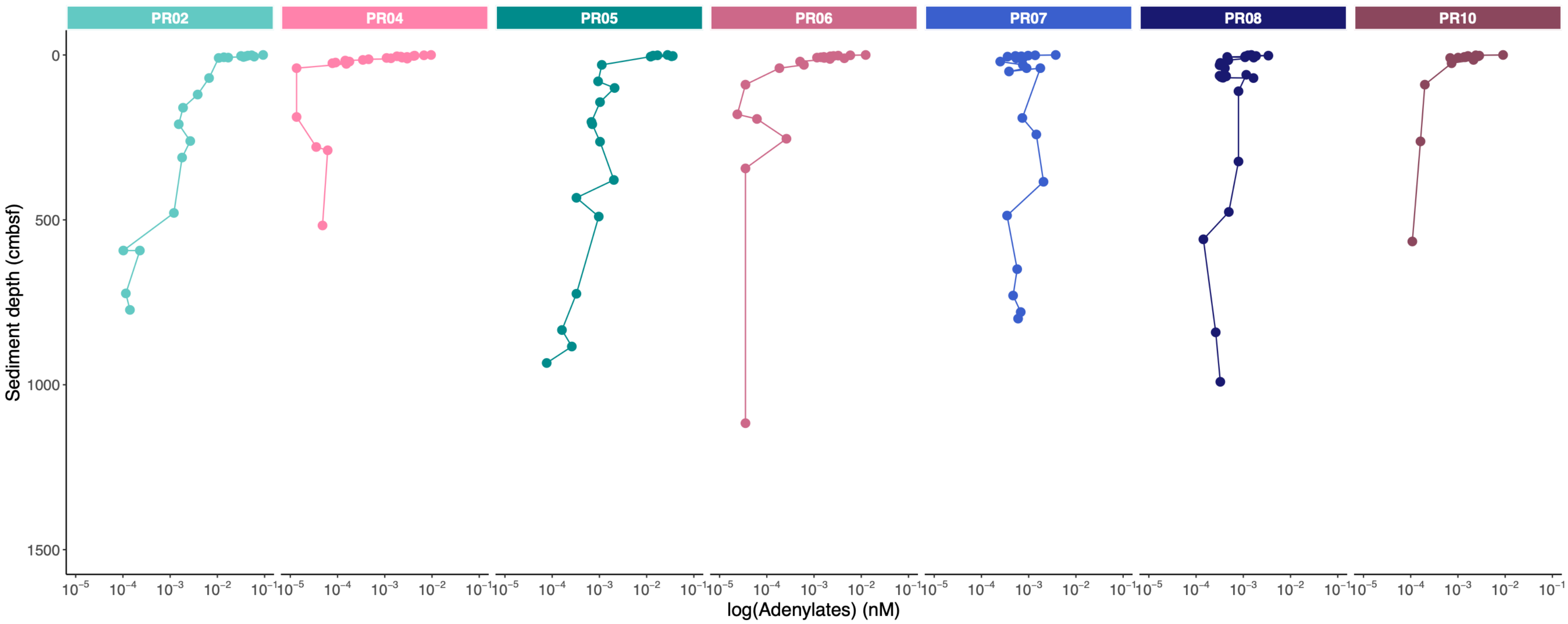


**Fig. S9** Profiles of sedimentary adenylate concentrations.

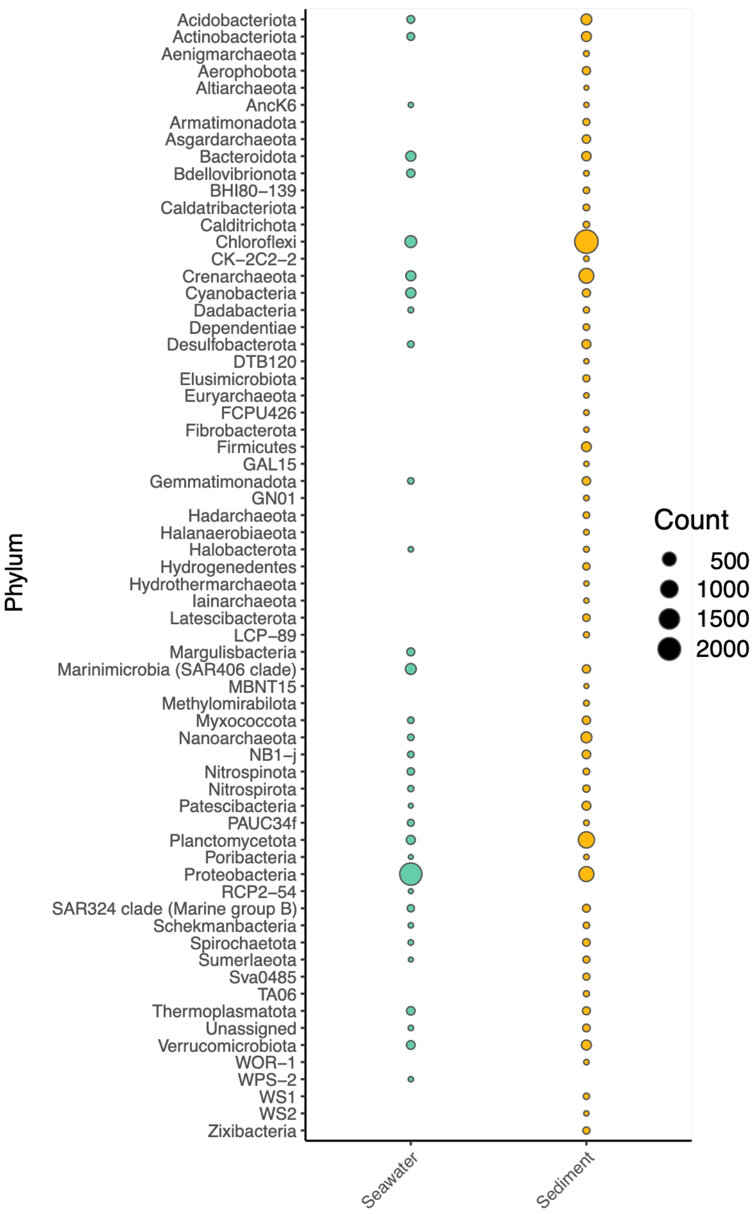


**Fig. S10** Differences in microbiome composition between seawater and sediment samples based on 16S rRNA amplicon analyses. Seawater and sedimentary communities show enrichment in 3,670 and 6,666 ASVs, respectively, most of which are members of the microbial phyla Proteobacteria, Chloroflexi, Planctomycetota and Crenarchaeota.

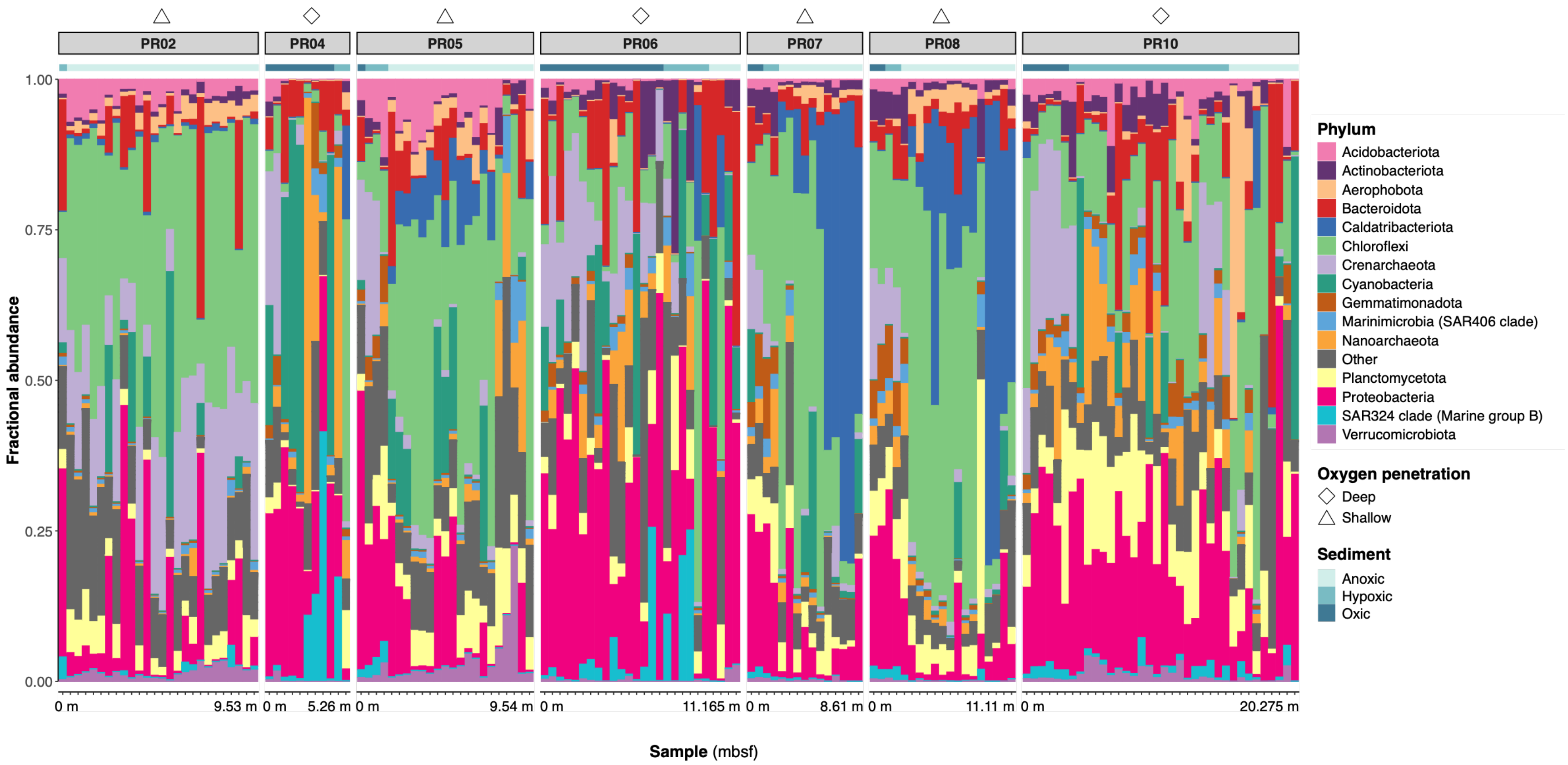


**Fig. S11** Relative abundances of microbial phyla in marine sediments.

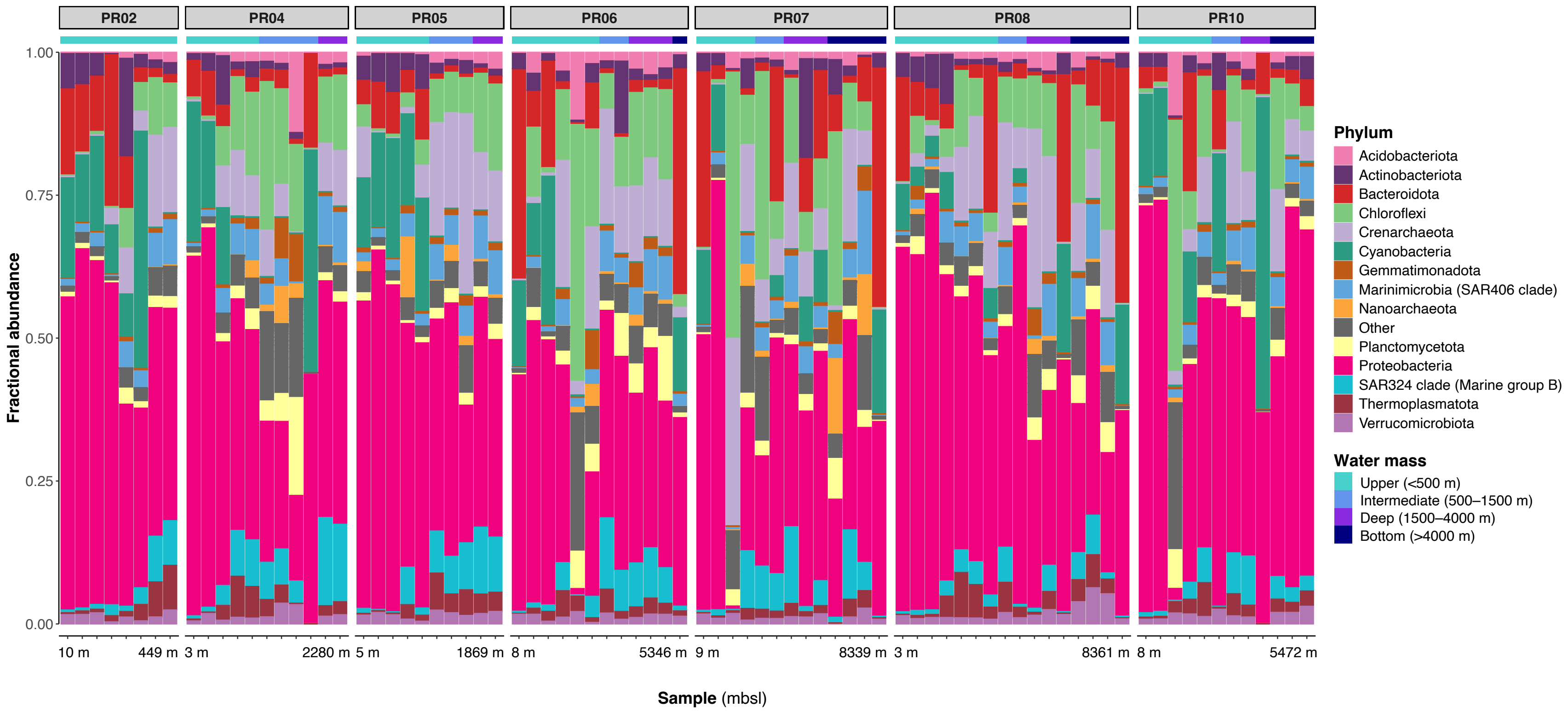


**Fig. S12** Relative abundances of microbial phyla in seawater.

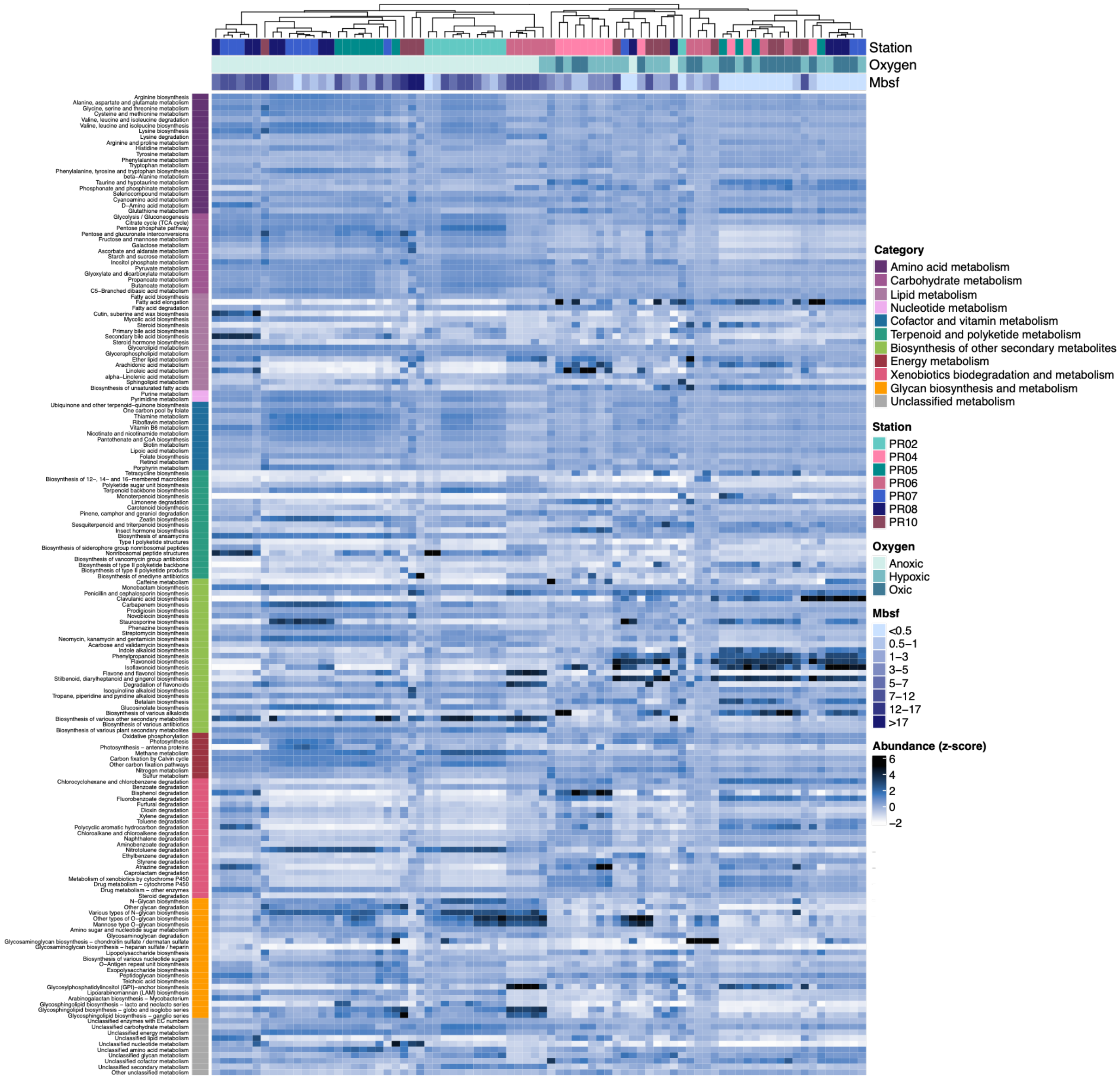


**Fig. S13** Full heatmap of average normalized read abundances assigned to KEGG metabolic pathways. Normalized read counts were summed for the corresponding metabolic pathways based on KO term identities and then divided by the number of genes assigned. The heatmap was calculated based on z-score scaled Euclidean distances.
